# Riemannian geometry of functional connectivity matrices for multi-site attention-deficit/hyperactivity disorder data harmonization

**DOI:** 10.1101/2021.09.01.458579

**Authors:** Guillem Simeon, Gemma Piella, Oscar Camara, Deborah Pareto

## Abstract

The use of multi-site datasets in neuroimaging provides neuroscientists with more statistical power to perform their analyses. However, it has been shown that imaging-site introduces a variability in the data that cannot be attributed to biological sources. In this work, we show that functional connectivity matrices derived from resting-state multi-site data contain a significant imaging-site bias. To this aim, we exploited the fact that functional connectivity matrices belong to the manifold of symmetric positive-definite matrices, making possible to operate on them with Riemannian geometry. We hereby propose a geometry-aware harmonization approach, Rigid Log-Euclidean Translation, that accounts for this site bias. Moreover, we adapted other Riemannian-geometric methods designed for other domain adaptation tasks, and compared them to our proposal. Based on our results, Rigid Log-Euclidean Translation of multi-site functional connectivity matrices seems to be among the studied methods the most suitable one in a clinical setting. This represents an advance with respect to previous functional connectivity data harmonization approaches, which do not respect the geometric constraints imposed by the underlying structure of the manifold. In particular, when applying our proposed method on data from the ADHD-200 dataset, a multi-site dataset built for the study of attention-deficit/hyperactivity disorder, we obtained results that display a remarkable correlation with established pathophysiological findings and, therefore, represent a substantial improvement when compared to the non-harmonization analysis. Thus, we present evidence supporting that harmonization should be extended to other functional neuroimaging datasets, and provide a simple geometric method to address it.

## 1 INTRODUCTION

Functional magnetic resonance imaging (fMRI) has become one of the leading methods to conduct research on human brain mapping. fMRI acquisitions allow the discovery of brain activation patterns, which help to understand the brain processes behind cognition or task performance. A relevant paradigm of acquisition is called resting-state fMRI, where the subject’s brain is studied without performing any specific task. Resting-state fMRI made possible to identify intrinsic functional connectivity and resting-state networks in the brain (van den Heuvel and Pol, 2010; Smitha et al., 2017), that is, distant brain regions that exhibit a temporal correlation in their blood-oxygen-level-dependent (BOLD) signals, and are thought to activate in a synchronous way. Moreover, resting-state functional connectomics has provided new insights into brain organization in disease states, since specific changes in connectivity patterns have been directly correlated to multiple disorders (Greicius, 2008; Du et al., 2018). These alterations in connectivity are useful to identify biomarkers or to gain more knowledge on these neurological or psychiatric disorders. A very informative approach summarizing whole-brain functional connectivity consists on the construction of functional connectivity matrices. It is based on the parcellation of the brain into some predefined regions of interest (ROIs) and the comparison of the BOLD time-series associated to these ROIs. Usually, this comparison is made in terms of a pair-wise Pearson’s correlation coefficient between ROI signals (even though partial correlations can also be used (Kim et al., 2015)). After the construction of functional connectivity matrices from a collection of subjects, one can identify entry-wise statistically significant differences between subsamples of these subjects (for example, patients and healthy controls), usually by performing univariate statistical tests on each entry separately. Therefore, one is able to spot key functional connections that display differences according to the condition of the subject and, hopefully, playing an important role in the disease state.

The fact that functional connectivity matrices are constructed using correlation coefficients makes them belong to a particular subset of matrices called symmetric positive-definite (SPD) matrices, which are symmetric matrices with all eigenvalues strictly greater than zero. This set of matrices does not form a vector space. This property implies that some conventional Euclidean operations on SPD matrices, such as subtraction of two SPD matrices, do not yield another element of the set of SPD matrices. Even though one can perform Euclidean operations with them, resulting matrices do not take into account the underlying geometry of the space of SPD matrices. Mathematically, the space of SPD is what is known as a manifold. Roughly speaking, a manifold is a curved space that locally looks like flat (Euclidean) space. Although one cannot perform Euclidean operations with elements (points) of the space, there exists an extremely powerful mathematical formalism, known as Riemannian geometry, that deals with manifolds and allows computations respecting their underlying geometry. Since functional connectivity matrices are SPD matrices, it is preferable to perform operations on them that take into account their underlying geometry, rather than applying standard Euclidean methods (You and Park, 2021). Geometry-aware approaches allow, for example, substantial increases in classification performance of functional connectivity matrices and accurate comparison of connectomes (Ng et al., 2014; Dodero et al., 2015; Varoquaux et al., 2010; Slavakis et al., 2017). Nevertheless, the potential of Riemannian geometric methods has not been fully recognized yet, though their consideration in functional studies is increasing. One remarkable contribution for the spread of these methods has been recently made by You and Park (You and Park, 2021), where they introduced SPDtoolbox, a MATLAB-based toolbox that offers the possibility of exploiting the geometry of functional connectivity matrices in a straightforward manner. One very relevant contribution put forward in (You and Park, 2021) is the proposal of a geometry-aware permutation testing framework that allows the identification of statistically significant differences in functional connectivity between groups.

Given the need for large cohorts to carry out statistical studies, together with the advent of big data and machine learning, neuroimaging datasets have increased their size usually by collecting data acquired at different sites. It is known that the use of different scanners, acquisition protocols or processing pipelines introduces a variability in the data that cannot be attributed to the biological variability of the subjects. To overcome this issue, the standardization of acquisition and processing protocols plays a key role. Nevertheless, it has been shown that some imaging-site bias remains in the data even when standardizing protocols and pipelines (David et al., 2013). Furthermore, from a machine learning perspective, classification might be more difficult because unwanted variability prevents the algorithm from learning the adequate hypothesis function, and also because the learned biased hypothesis might not have the necessary generalization power to unseen examples, affecting classification performance (Brain and Webb, 2002). One well-known approach to harmonize neuroimaging data is ComBat (Johnson et al., 2006). Essentially, ComBat involves subtracting batch variability by modelling it as a deviation from the estimated influence of known covariates of the model. By taking into account the effect of covariates, one can control for the other sources of variability other than the batch (site) effect. In the field of neuroimaging, the ComBat method was shown to remove imaging-site effects in diffusion tensor imaging acquisitions and cortical thickness studies (Fortin et al., 2017, 2018; Beer et al., 2020). Later, ComBat was adapted to functional acquisitions and their derived functional connectivity matrices (Yu et al., 2018).

Nevertheless, this approach does not take into account the geometry of the space of SPD matrices and, therefore, it cannot be considered to be operating in a geometry-constrained manner. In fact, the proposed adaptation of ComBat to functional connectivity matrices (Yu et al., 2018) only considers that resulting harmonized matrices need to be symmetric. Nevertheless, positive-definiteness is not enforced and, therefore, resulting matrices do not belong in general to the original space of SPD matrices. Positive-definiteness is a geometric constraint and precisely defines the manifold of SPD matrices as a subspace of the space of symmetric matrices. Any method that operates on SPD matrices and respects their symmetry and positive-definiteness can be regarded as a geometry-constrained method. Although site harmonization using Riemannian methods has not been directly addressed, some proposals have been put forward to deal with related procedures. In particular, there have been previous works on the use of manifold-constrained operations to develop domain adaptation procedures specifically for functional data (Ng et al., 2014; Yair et al., 2019). However, in their case, domain adaptation was performed targeting inter-session variability (that is, differences arising from acquiring data from the same subject in different sessions, possibly with intervention in-between).

Our intention in this work is to use the rich formalism of Riemannian geometry to geometrically characterize imaging-site bias and to propose methods to remove it (at least partially) from multi-site functional connectivity matrices. To this aim, we will adapt two previous contributions to domain adaptation of functional data and we will also propose a method from our own, called Rigid Log-Euclidean Translation. These operations will be particularized to ADHD-200 (ADHD-200 Consortium, 2017), a multi-site dataset of resting-state acquisitions from attention-deficit/hyperactivity (ADHD) patients and healthy controls (HC). Furthermore, using the geometry-aware permutation testing algorithm proposed in (You and Park, 2021), implemented in SPDtoolbox, we will analyse the impact of imaging-site bias on the results that can be distilled from functional connectivity matrices, and we will correlate them with established pathophysiological findings in ADHD. In short, we intend to provide a simple but powerful geometrically-grounded framework for multi-site functional connectivity matrices harmonization.

## 2 MATERIAL AND METHODS

### 2.1 Functional connectivity matrices from ADHD-200

The ADHD-200 sample (ADHD-200 Consortium, 2017) is the result of a joint international effort that provides researchers worldwide with openly-shared resting-state and structural acquisitions from 8 different sites, comprising 362 children and adolescents diagnosed with ADHD and 585 typically developing controls, along with their phenotypic data (ADHD subtype, ADHD score, IQ, medication status, etc). In an effort to standardize the analyses carried out with ADHD-200, three different preprocessing pipelines for resting-state acquisitions were proposed, being also preprocessed data made publicly available (Bellec et al., 2017). One of this preprocessing strategies, the Athena pipeline, was performed by Craddock (Craddock, 2011). The relevant derived data for our present work are the extracted ROI time-courses from resting-state acquisitions. As previously mentioned, the computation of the Pearson’s correlation coefficient between time-courses allows the construction of functional connectivity matrices. Craddock provided researchers with time-series extracted using different parcellation schemes. In particular, Craddock derived specific functional parcellations for ADHD-200 data from their resting-state acquisitions, giving rise to a parcellation into 190 ROIs (CC200).

For our present work, we have used a subset of functional connectivity matrices constructed using the CC200 parcellation scheme. Specifically, our dataset has consisted of functional connectivity matrices from 80 ADHD patients and 80 HC. Their acquisitions have been randomly pooled (avoiding any kind of matching in order to maximize intrinsic variability) from four different sources: 20 ADHD/20 HC from Kennedy Krieger Institute (KKI) (Johns Hopkins University), 20 ADHD/20 HC from the NeuroIMAGE sample (NIM), 20 ADHD/20 HC from New York University Child Study Center (NYU), and again 20 ADHD/20 HC from Peking University (PKG). Therefore, we have worked in total with 160 functional connectivity matrices, with dimensions of 190 × 190.

### 2.2 Riemannian geometry of SPD matrices

Symmetric positive definite (SPD) matrices are the set of symmetric matrices with all eigenvalues strictly greater than zero. This property is equivalent to the requirement that, for a *n* × *n* symmetric matrix Σ to be positive definite, *x*^*T*^ Σ *x >* 0 for all non-zero *x* ∈ ℝ^*n*^. The set of SPD matrices forms a manifold, a topological space that locally looks like Euclidean space. At each point *p* of some manifold ℳ, one can construct a tangent space 𝒯 *_p_ ℳ*, generated by the tangent vectors of curves crossing *p*, which is a vector space and approximates the manifold in the neighbourhood of *p*. In our present case, ℳ is the manifold of *n* × *n* SPD matrices, which we will denote by 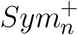, where each point is a SPD matrix, and one can show that at every point 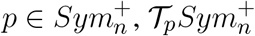 is the space of symmetric *n* × *n* matrices, *Sym*_*n*_.

Riemannian geometry provides a framework that allows one to regard the manifold as a metric space. Specifically, a Riemannian manifold is a smooth manifold equipped with a symmetric, positive-definite tensor field called metric tensor. The metric tensor *g* at every point *p* ∈ ℳ is a map

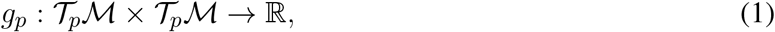

constructed as a generalization of the canonical dot product in Euclidean space, and that can be used to characterize the geometry of the manifold. Two other important maps in Riemannian geometry are the exponential map exp_*p*_ and the logarithmic map log_*p*_ at some point *p* ∈ ℳ,

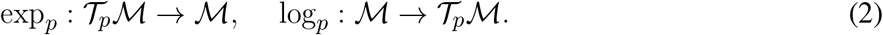

Given a tangent vector *v* ∈ 𝒯_*p*_ℳ, exp_*p*_(*v*) results in a projection of *v* onto the neighbourhood of *p* in the manifold ℳ by defining a geodesic (i.e., a shortest length curve on the manifold) in the direction of *v*. The logarithmic map performs the inverse operation, mapping points in the manifold to vectors in the tangent space of *p*.

Particularizing previous notions to our case, Pennec et al. (Pennec et al., 2006) introduced one of the most used geometric Riemannian structures on the manifold of SPD matrices, 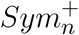, the affine-invariant Riemannian metric (AIRM). They proposed the following metric tensor at some point 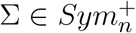 (a SPD matrix) applied on two tangent vectors *X, Y* ∈ *Sym*_*n*_ (two symmetric matrices):

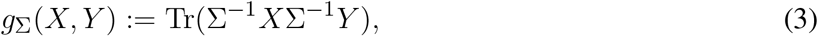

where Σ^−1^ is the matrix inverse and Tr(·) is the usual trace operator. Associated to this metric, the exponential map at Σ acting on a tangent vector *V* ∈ *Sym*_*n*_ and the logarithmic map at Σ acting on a point 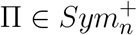 read

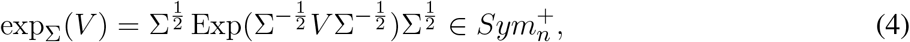

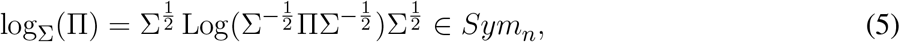

being Exp(*A*) and Log(*A*) the exponential and logarithm of matrix *A*, respectively, which can be computed after eigenvalue decomposition *A* = *UDU*^*T*^ as Exp(*A*) = *U* exp(*D*) *U*^*T*^, Log(*A*) = *U* log(*D*) *U*^*T*^, where *U* is the eigenvector matrix and *D* the diagonal eigenvalue matrix, and exp(*D*) (log(*D*)) denotes the application of the exponential (logarithmic) function to the eigenvalues. 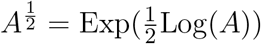 is the unique SPD square root of the SPD matrix *A*. Finally, under the AIRM framework, the geodesic distance between two SPD matrices Σ and Π is

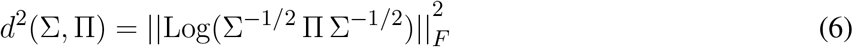

With 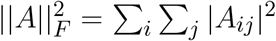, the (squared) Frobenius norm of a matrix *A*. Arsigny et al. (Arsigny et al., 2005) proposed another framework to account for the Riemannian structure of the manifold of SPD matrices. This framework, known as Log-Euclidean Riemannian Metric (LERM), consists basically on embedding points of the manifold in an Euclidean space by using the matrix logarithm Log(·). Standard Euclidean computations can be performed on SPD matrices’ logarithms, since they have been mapped to Euclidean space. Under LERM, the geodesic distance between two SPD matrices Σ, Π is simply

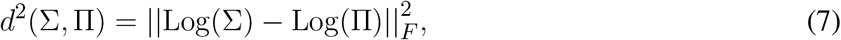

that is, the Euclidean distance of matrix logarithms. The computation of these distances is directly implemented in SPDtoolbox.

A useful object in the manifold that can be constructed given a collection of SPD matrices Σ_1_, Σ_2_, …, Σ_*N*_ is their Fréchet mean. The Fréchet mean is the generalization of the concept of centroid or center of mass to more general metric spaces other than Euclidean space, giving a sense of centrality measure. Mathematically, the Fréchet mean of the previous set of matrices is defined as

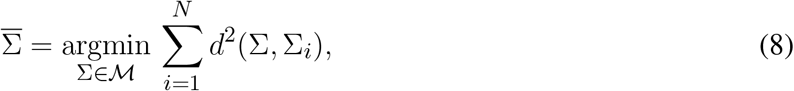

that is, the SPD matrix 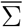 that minimizes the sum of the squared distances from itself to the collection of matrices. The Fréchet mean depends on the chosen distance function in the manifold and, therefore, the Fréchet means under LERM and AIRM in general do not coincide. When considering AIRM, the Fréchet mean has to be computed by an optimization procedure. In (Pennec, 2006), the author proposed a Newton gradient descent algorithm which, after some mean initialization, consists on iterating the following update rule to get successive estimates 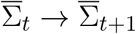 until convergence:

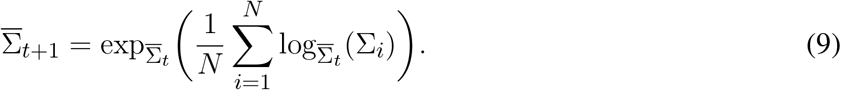

The algorithm involves the computation of several matrix logarithms, square roots, inverses and exponentials at each iteration, resulting in a computationally expensive process, especially when dealing with a large number of samples. On the contrary, under LERM, the Fréchet mean has a closed form that enables a very fast computation:

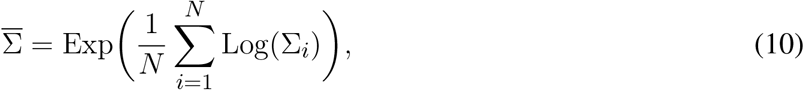

which reduces to the obtention of the Euclidean mean of the matrices’ logarithms (vectors) and its projection back to the manifold via exponential map. Again, both Fréchet mean computation methods are directly implemented in SPDtoolbox.

### 2.3 Geometry-aware permutation testing

Permutation tests are non-parametric statistical tests that rely on the randomization of the observed data to assess the statistical significance of group differences (Nichols and Holmes, 2001). In our case, we have collected two sets of functional connectivity matrices, one with *N*_+_ ADHD patients 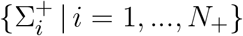 and the other one with *N*_−_ healthy controls 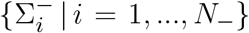. The philosophy behind permutation testing is to make a very large number of random permutations of the *N* subjects into the two subgroups and to sample the distribution of values of some statistic under the null hypothesis, that is, under the assumption that the allocation of a subject into one of the two groups is arbitrary. This procedure allows the obtention of the approximate null distribution of the statistic and the assessment of how extreme this observation is with respect to the null distribution of values. Specifically, one can use the definition of p-value to reject the null hypothesis, i.e., the probability to obtain a value for the statistic as or more extreme than the one observed, under the assumption that the null hypothesis is correct. Therefore, under the permutation testing approach, the p-value is the proportion of sampled permutations where the statistic is greater than or equal to the originally observed statistic.

The geometry-aware algorithm proposed in (You and Park, 2021) and implemented in SPDtoolbox has the following steps:

1. Compute Fréchet means 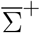 and 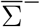 ADHD patients and healthy controls, respectively, and take 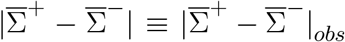 (the entry-wise absolute value of the difference matrix, the observed statistic of the *n* × *n* entries).
2. For *T* iterations, perform permutations of the subjects of sizes *N*_+_ and *N*_−_ as previously described, recompute group Fréchet means, and record the value of the statistic at iteration 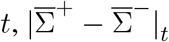.
3. For each entry *ij*, compute its corresponding p-value by taking the ratio between the number of iterations giving a statistic greater than or equal to 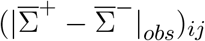 and the total number of iterations *T*.

At this point, assuming some confidence level *α*, one can reject the null hypothesis for entries with p-values lower than *α*. It is worth noting that the algorithm tests for statistical significance in absolute difference, and therefore cannot provide information on the positive or negative character of the difference. The output of the algorithm in our case will be a 190×190 matrix with p-values as entries. Nevertheless, we will not rely on a single permutation test on the 80 ADHD/80 HC subjects to perform our analyses. Instead, we have decided to perform 100 separate permutation tests on 10 ADHD/10 HC randomly sampled subjects in each experiment. The reasons for following this approach are: first, taking into account the philosophy behind permutation testing, performing a reliable permutation test on 80 ADHD and 80 HC is not computationally tractable, since the number of ways of dividing 160 subjects into two separate groups of 80 is 160!*/*(80! 80!) ∼10^47^ and, therefore, by no means one can sample a large enough fraction of permutations to rely on the obtained distribution of statistic values; second, by randomly pooling 10 ADHD/10 HC 100 times, regardless of imaging site, one can probe different combinations of site members and site influence on the variability of the results; and third, one can assess the relevance of salient connections and nodes by checking for coincidences between independent permutation tests. Taken together, this treatment allows evaluating the power of harmonization methods, in the sense that when data is harmonized, one expects more sensitivity (coherence, coincidences) in statistically significant results when using randomly sampled subjects from different sites. Given these aforementioned points, we have operated in the following manner: we have performed 100 separate permutation tests, each one of 1000 iterations, on 10 ADHD/10 HC randomly sampled subjects; after each test, we have identified the entries with *p <* 0.001 and declared them as significant; at this point, we have been left with 100 binary matrices of dimensions 190 ×190 (one for each separate test) with ones in the location of significant entries and zeros elsewhere; and, finally, we have summed these binary matrices to obtain a symmetric 190 ×190 matrix, which we call *F*, with *F*_*ij*_ being the number of times the connection *ij* has been declared significant, that could be regarded as a 2-dimensional histogram.

When working at a significance level *α* = 0.001 for each individual permutation test, we are assuming that, at most, the probability of mistakenly declaring an entry as statistically significant by chance is 0.001 (incorrect rejection of the null hypothesis). However, when performing 100 separate experiments and considering jointly their results (as in the frequency histograms), this effect can be accumulated. As a simplification, we can model each separate experiment as a random Bernoulli experiment: the probability of mistakenly declaring each entry as significant by chance in a separate experiment is 0.001. The concatenation of several independent Bernoulli experiments follows the binomial distribution. Therefore, we can use the binomial distribution to compute the probabilities for each separate entry to be declared significant by chance once, twice, 3 times, etc. In our case, considering a probability of 0.001 and 100 trials, the binomial distribution predicts that in the final frequency matrix, the probability for each entry to appear once is 0.09. That is, each entry has a probability of 9% of appearing once in the frequency histogram by chance, what indeed means that at this frequency level (*F*_*ij*_ = 1), significance is compromised. However, for entries *F*_*ij*_ ≥ 2, the (complementary cumulative) probability is ≈ 0.005, and for entries *F*_*ij*_ ≥ 3 the resulting (complementary cumulative) probability is ≈ 0.00015. As a consequence, one can threshold the histogram matrix *F*_*ij*_ at different frequencies depending on the confidence that wants to assume.

Finally, using these notions and taking into account that the previously described procedure will be applied to different imaging-site harmonization methods introduced in subsection 2.4, we define a simple quantity that measures the power of a method to specifically point to some entries by accumulating counts across the 100 separate experiments. We call this measure the sensitivity of the method. At a given frequency *n*, the sensitivity *S*(*n*) was defined as the ratio between the number of entries such that *F*_*ij*_ ≥ *n* and the number of entries such that *F*_*ij*_ ≥ 1.

### 2.4 Geometry-aware site harmonization

As mentioned in the Introduction, imaging site has an impact on the acquisition of functional data and their analyses. We show that this is indeed the case for ADHD-200, even when using the same preprocessing pipeline (Wang et al., 2017). We also prove that, in our case, the principal difference stems from the biased and site-clustered distribution of connectivity matrices in the SPD manifold. Some geometry-aware domain adaptation approaches have been proposed, even though not targeting imaging-site variability. We can distinguish two different harmonization methods that will be analysed separately: matrix whitening at identity and parallel transport. When being described, they will be adapted to our dataset of ADHD patients 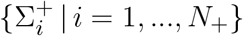 and healthy controls 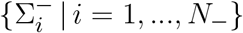 and to our imaging-site harmonization aim.

#### 2.4.1 Matrix whitening at identity

Suppose our dataset can also be divided into collections of matrices 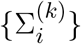, with *k* labeling the imaging site where resting-state data from subject *i* was acquired. If we denote by 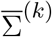 the AIRM Fréchet mean of matrices obtained from site *k*, the matrix whitening approach reduces to applying the following transformation to all 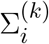 (Ng et al., 2014; Yair et al., 2019):

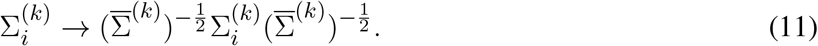

One can show that this transformation is equivalent to a displacement of the matrices such that their Fréchet mean is the identity matrix. To this aim, consider *C* ∈*GL*(*n*), that is, an invertible *n* × *n* matrix. The following properties hold when considering the AIRM framework (Yair et al., 2019):

1. The geodesic distance (6) between two SPD matrices Σ and Π is invariant under Σ → *C* Σ*C*^*T*^ and Π → *C* Π*C*^*T*^, i.e., *d*(*C* Σ*C*^*T*^, *C* Π*C*^*T*^) = *d*(Σ, Π).
2. Given some Fréchet mean 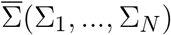, then 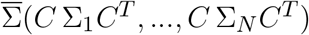 is equivalent to 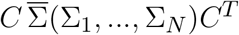.

Transformation (11) can be expressed as 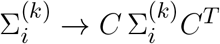 with 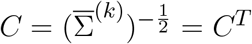, following the last equality from the fact that, if 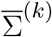 is SPD, 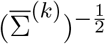 is also SPD and, therefore, 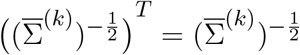. By virtue of the first property, intra-site geodesic distances are preserved, and the Fréchet mean becomes 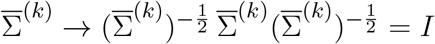, following from the second property.

In (Ng et al., 2014), authors followed this approach and applied the logarithmic map Log(·) to project transformed SPD matrices to the common tangent space at identity before performing classification. To our understanding, invariance of intra-site distances is a desirable property for harmonization since it is essential to preserve intra-site variability, presumably coming from biological sources (that is, disease, age, gender, medication status, etc). However, we think that, in our setting, it might be a priori more cautious not to remove information about the original location of matrices in the manifold by displacing them to the neighbourhood of the identity matrix, since it could also remove relevant clinical information.

One could also consider the use of two different 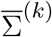, one for patients and another one for healthy controls, since one expects their Fréchet means to be different. However, the use of a single site Fréchet mean to perform the transformation is based on two considerations. On the one hand, the use of different transformations for ADHD patients and HC would violate the preservation of all pair-wise distances within the site, since the matrix *C* would be different depending on the condition of the subjects. Distances among HC on one side and distances among ADHD patients on the other side would be preserved, while distances between HC and ADHD patients would not. This fact could be regarded as the introduction of an uncontrolled source of variability between conditions. On the other hand, it would be desirable to have a harmonization approach that does not depend on the availability of knowledge about subjects’ conditions, which would be required when using different transformations. For example, as stated in the Introduction, domain adaptation has been proven useful when training classifiers. Ideally, when the classifier is reliable enough, one would like to assign the condition to a subject without knowing it a priori. Therefore, the use of a single transformation would allow the harmonization of the data prior to the use of the classifier.

#### 2.4.2 Parallel transport

Elements from two different tangent spaces cannot be directly compared. To compare them in an appropriate manner, one needs to perform what is called parallel transport. In short, parallel transport refers to transporting a tangent space at a point along a geodesic distorting vectors of this tangent space as less as possible. In (Yair et al., 2019), authors provide an excellent and rigorous mathematical description of parallel transport in the manifold of SPD matrices. For harmonization and comparison of SPD matrices, we are interested in transporting and projecting them to a common tangent space.

The main line of reasoning applied to our case is as follows:

1. Project matrices from a given site *k* to the tangent space of their site Fréchet mean 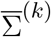 via logarithmic map 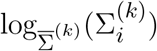.
2. Parallel transport them to the tangent space of a common reference point Σ_0_.
3. Reproject the parallel transported matrices back to the manifold via exponential map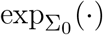.

One of the main results of (Yair et al., 2019), is that all this process can be performed by the transformation 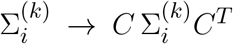 with 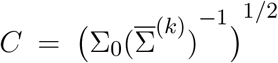. Therefore, properties 1 and 2 presented in the case of matrix whitening also hold here: intra-site distances are preserved and the Fréchet mean becomes 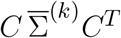.

Then, one has to decide which is the appropriate reference point Σ_0_. Ng et al. (Ng et al., 2014) used Σ_0_ = *I* to compare the classification performance between the present method and matrix whitening at identity, although they performed parallel transport numerically (Schild’s ladder). In this case, 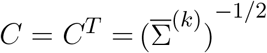 and the Fréchet mean reduces to *I* (notice that the transformation is exactly equivalent to matrix whitening), arising the same previously mentioned concerns about original location in the manifold. On the other hand, Yair et al. (Yair et al., 2019) suggest using as the reference point 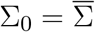, the Fréchet mean of site means 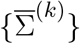. This approach takes into account the original location of SPD matrices in the manifold through the global mean 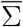. Right after this step, Yair et al. perform matrix whitening using 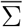. We will refer to this whole process proposed by Yair et al. as simply parallel transport (even though proper parallel transport is the step performed after projecting to the site mean’s tangent space and before reprojecting back to the manifold at the reference point). Notice that the use of a single reference point Σ_0_ in our setting, both for ADHD patients and HC, can be justified by taking into account the same considerations from the previous subsection.

A priori, matrix whitening and parallel transport are different transformations. Nevertheless, one can show that under the condition 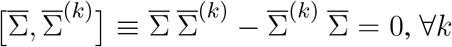, that is, when site means commute with the global mean, parallel transport and matrix whitening are equivalent frameworks. This property is proven in the Annex, and it will have an impact on the results we obtain.

#### 2.4.3 Rigid Log-Euclidean Translation

We hereby propose a method for site harmonization, which we call Rigid Log-Euclidean Translation (RLET), that has the two a priori desirable properties stated above: preservation of intra-site geodesic distances, thus retaining biological variability, and consideration of the original location of correlation matrices in the manifold. Site bias could also impact the distribution of matrices around site mean, but this would require modelling site effects on the distribution of matrices around their mean. The proposed method is therefore a geometrically-motivated first order approximation. We will work under the LERM framework, which allows a purely Euclidean treatment of the logarithm of the matrices. The steps of Rigid Log-Euclidean Translation are the following ones:

1. Compute the logarithm of site Fréchet mean for all *k*

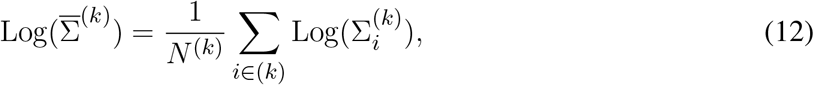

where the sum runs through all subjects in site *k*, and *N* ^(*k*)^ is the total number of subjects in site *k*.
2. Compute the logarithm of the global mean 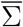 defined as

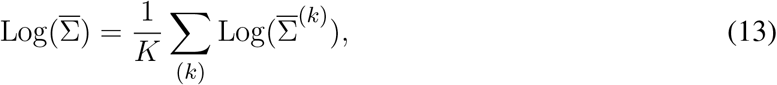

with *K* being the number of sites.
3. Apply the following transformation to the entire collection of matrices 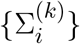 given their *k*

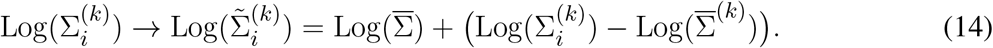
4. Use the exponential map to obtain the modified (site-harmonized) SPD matrices 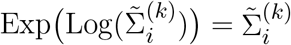.

When taking into account that after applying the logarithmic map to the original matrices we obtain vectors of the Euclidean space ℝ^*n*×*n*^, the above transformation is merely a translation of the original matrices such that transformed site means 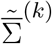 are identically the global mean 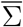. In fact, it is a rigid translation of matrices logarithms, representing an isometry of the Euclidean metric, and therefore preserving Euclidean distances in the Euclidean space ℝ^*n*×*n*^. These properties are formally proven in the Annex.

Since geodesic distances under the LERM framework are these Euclidean distances, intra-site geodesic distances are invariant under the proposed transformation, and the Rigid Log-Euclidean Translation approach preserves intra-site variability. The transformation also allows retaining information about the original location in the manifold through the global mean. On the other hand, one can straightforwardly modify this approach to rigidly transport matrices to *I*, by removing the term 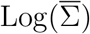 from the transformation rule. In this case 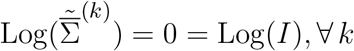, (see Annex) and resulting relative positions between all matrices in Euclidean space are the same as when translated to 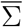. We will refer to these two approaches as 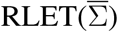 and RLET(*I*). RLET(*I*) can be useful to compare different harmonization strategies at identity, and to probe differences coming uniquely from manifold location.

### 2.5 Pair-wise distances and low-dimensional embedding

Even though all the concepts presented above are geometrically motivated, the major difficulty for bias characterization stems from the fact that we are dealing with a very high-dimensional space (in our case, *d* = *n*(*n* + 1)*/*2 = 190(190 + 1)*/*2 = 18145). Therefore, the extraction of insights from this space is a challenging task. However, since our manifold is equipped with a metric and a distance measure (i.e., the manifold is Riemannian), we can construct a useful object that can summarize important geometric information: the pair-wise distance matrix *D*_*ij*_ = *d*(Σ_*i*_, Σ_*j*_) between functional connectivity matrices (subjects). In fact, this approach of using pair-wise distances or pair-wise similarities is common in unsupervised techniques of dimensionality reduction for visualization purposes, such as multidimensional scaling (MDS) or t-distributed stochastic neighbour embedding (tSNE).

We will focus on t-distributed stochastic neighbour embedding (tSNE), since it was the method of choice in the previous works we have adapted for site harmonization. tSNE (van der Maaten and Hinton, 2008) is a nonlinear dimensionality reduction technique that maps a data point in a high-dimensional space to a point in 2-dimensional or 3-dimensional Euclidean space. Differently from other nonlinear dimensionality reduction approaches, tSNE has a probabilistic nature. Originally, the similarity metric of the algorithm was taken to be the pair-wise Euclidean distance between high-dimensional points, although currently different metrics can be used depending on the task at hand. However, the usual Euclidean distance is a good choice in our case: by using as high-dimensional inputs the logarithms of the matrices, the similarity measure between two different matrices will turn out to be the LERM geodesic distance. As a consequence, the low-dimensional embedding and its visualization will display similarity given by the pair-wise distribution of matrices in the manifold (under the LERM framework) with very high probability. At this point, one is able to assess the distribution of matrices in the manifold and, therefore, to characterize original geometric imaging-site bias and evaluate the effect of harmonization methods.

### 2.6 Functional analysis

One can perform comparative functional analyses between methods in terms of resting-state networks instead of specific ROI-to-ROI connections. To this aim, we will use the Yeo 17-networks atlas (Yeo et al., 2011), where the brain is parcellated into 17 different resting-state networks. We have further reduced the number of networks by using the following proposed correspondence: 1 and 2 correspond to the visual network (VIS); 3, 4 and 14 to the sensorimotor network (MOT); 5 and 6 to dorsal attention (DA); 7 to ventral attention (VA); 8, 11, 12 and 13 to frontoparietal network (FP); 9 and 10 to limbic (LIM); and 15, 16 and 17 to default mode network (DMN). However, Yeo’s parcellation does not include important subcortical regions that are included in our ROIs (CC200 parcellation, see Introduction). Therefore, and taking into account the results we have obtained, we have introduced the brain-stem (BS), the cerebellum (CRB) and basal ganglia (BG). In total, we are left with 10 different functional components.

Once we have the functional parcellation of the brain, we will assign each ROI to its corresponding functional component by using the ROI’s coordinates. This will allow us to initially focus our attention on inter-network and intra-network interactions, since the individual study of salient connections would be intractable when the number of detected significant differences in ADHD and HC connectomes is large. When thresholding frequency matrices *F*_*ij*_ at higher frequency levels, the number of samples will be reduced and an individual treatment of these differences will be possible.

## 3 RESULTS

### 3.1 Pair-wise distances and distribution of matrices

We have constructed the aforementioned pair-wise distance matrices *D*_*ij*_ = *d*(Σ_*i*_, Σ_*j*_) that retain information of the high-dimensional distribution of the matrices in the manifold. Specifically, we have computed pair-wise distances for: original (non-modified) matrices *D*_*ij*_^(0)^ both using AIRM and LERM distances, harmonized matrices under Rigid Log-Euclidean Translation 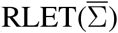 and RLET(*I*)), 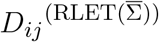 and *D*_*ij*_^(RLET(*I*))^, using LERM distance, and matrix-whitening-harmonized (MW) and parallel-transport-harmonized matrices (PT), *D*_*ij*_^(MW)^ and *D*_*ij*_^(PT)^, employing AIRM distance (Figure 1).

**Figure 1.**
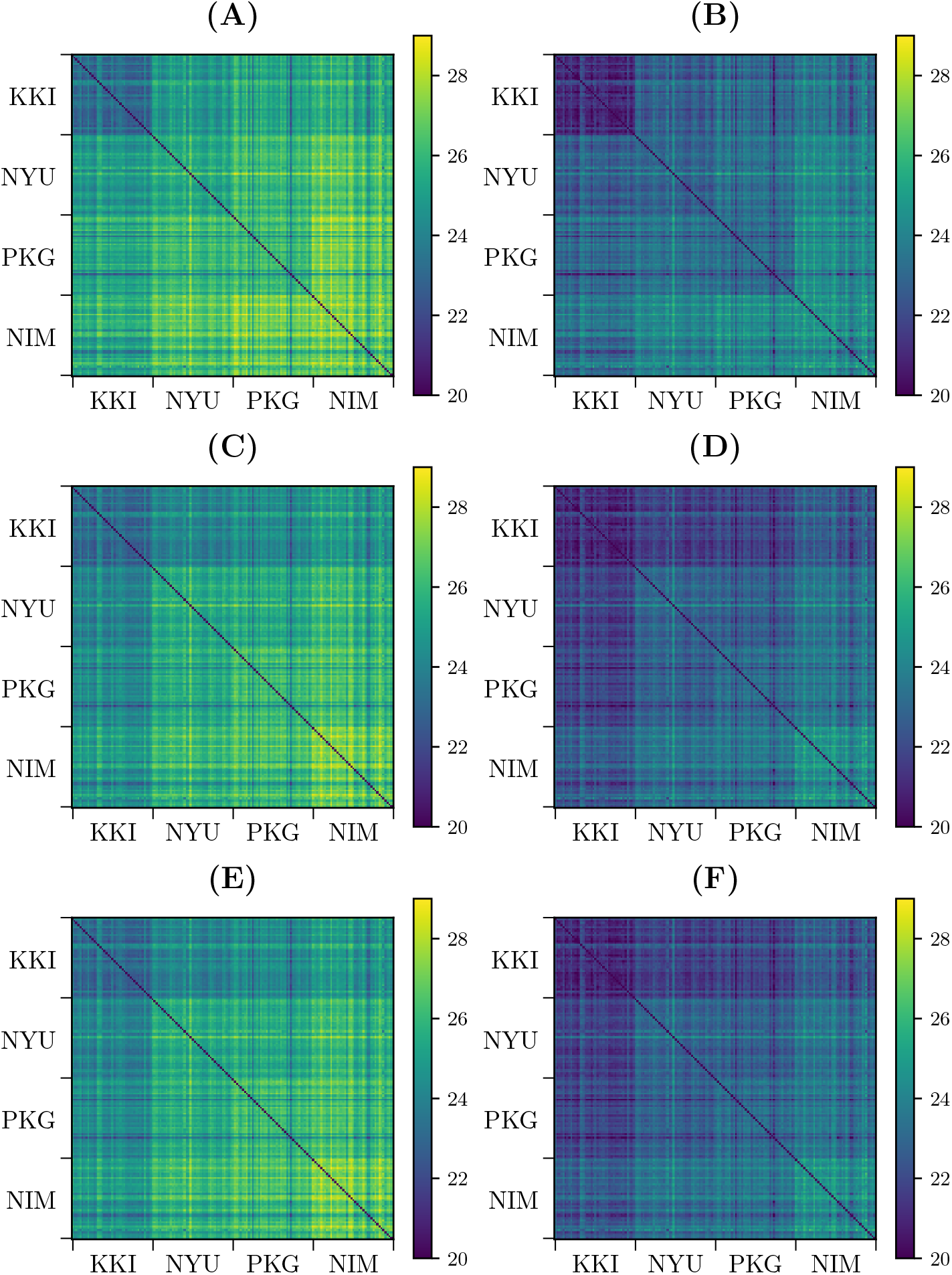
Pair-wise distances between functional connectivity matrices from different sites (KKI: Kennedy Krieger Institute, Johns Hopkins University; NYU: New York University Child Study Center; PKG: Peking University; NIM: NeuroIMAGE sample). **(A)** Pair-wise affine-invariant Riemannian metric (AIRM) distances for non-transformed matrices. **(B)** Pair-wise Log-Euclidean Riemannian metric (LERM) distances for non-transformed matrices. **(C)** Pair-wise AIRM distances after matrix whitening. **(D)** Pair-wise LERM distances after Rigid Log-Euclidean Translation (RLET) to the global mean 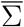. **(E)** Pair-wise AIRM distances after parallel transport. **(F)** Pair-wise LERM distances after RLET to the identity matrix.

*D*_*ij*_^(MW)^ and *D*_*ij*_^(PT)^ look identical (Figure 1C and Figure 1F). This approximate equivalence can be explained by nearly vanishing commutators 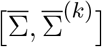, as previously proved. We have checked the values of these commutators and, indeed, they almost vanish, as can be seen in Figure 2. The consequence is that, in practical terms, in the following we can regard PT and MW as two equivalent frameworks.

**Figure 2.**
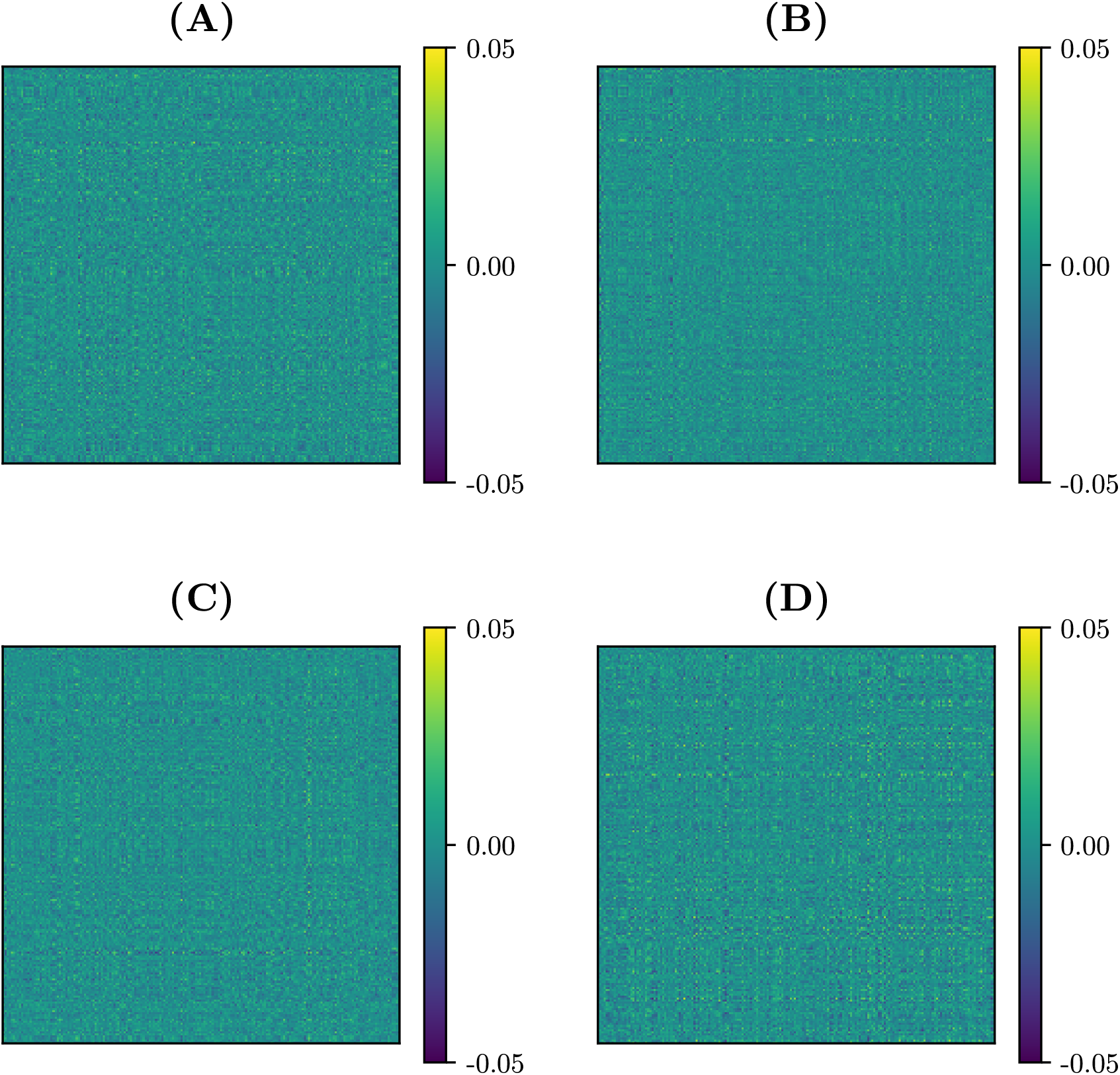
Commutators of the global mean 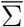 and original site Fréchet means 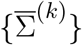. **(A)** 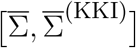 **(B)** 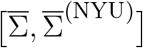 **(C)** 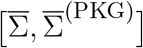 **(D)** 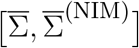. (KKI: Kennedy Krieger Institute, Johns Hopkins University; NYU: New York University Child Study Center; PKG: Peking University; NIM: NeuroIMAGE sample).

On the other hand, we have applied tSNE to the logarithms of: original (non-harmonized) matrices, 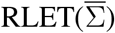-harmonized matrices and MW-harmonized matrices, with results shown in Figure 3 and Figure 4. Therefore, we have obtained a low-dimensional representation of 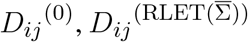 and *D*_*ij*_^(MW)^, all of them computed by means of LERM distances. RLET(*I*)-harmonized matrices have not been used since, by construction, their pair-wise distances are identical to the ones for 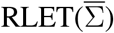, and, regarding PT-harmonized matrices, they are almost exactly equivalent to the ones obtained using MW. The differences that could be displayed in the visualization of RLET(*I*) matrices compared to 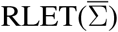-harmonized ones and PT compared to MW would be due to the stochastic nature of the tSNE algorithm.

**Figure 3.**
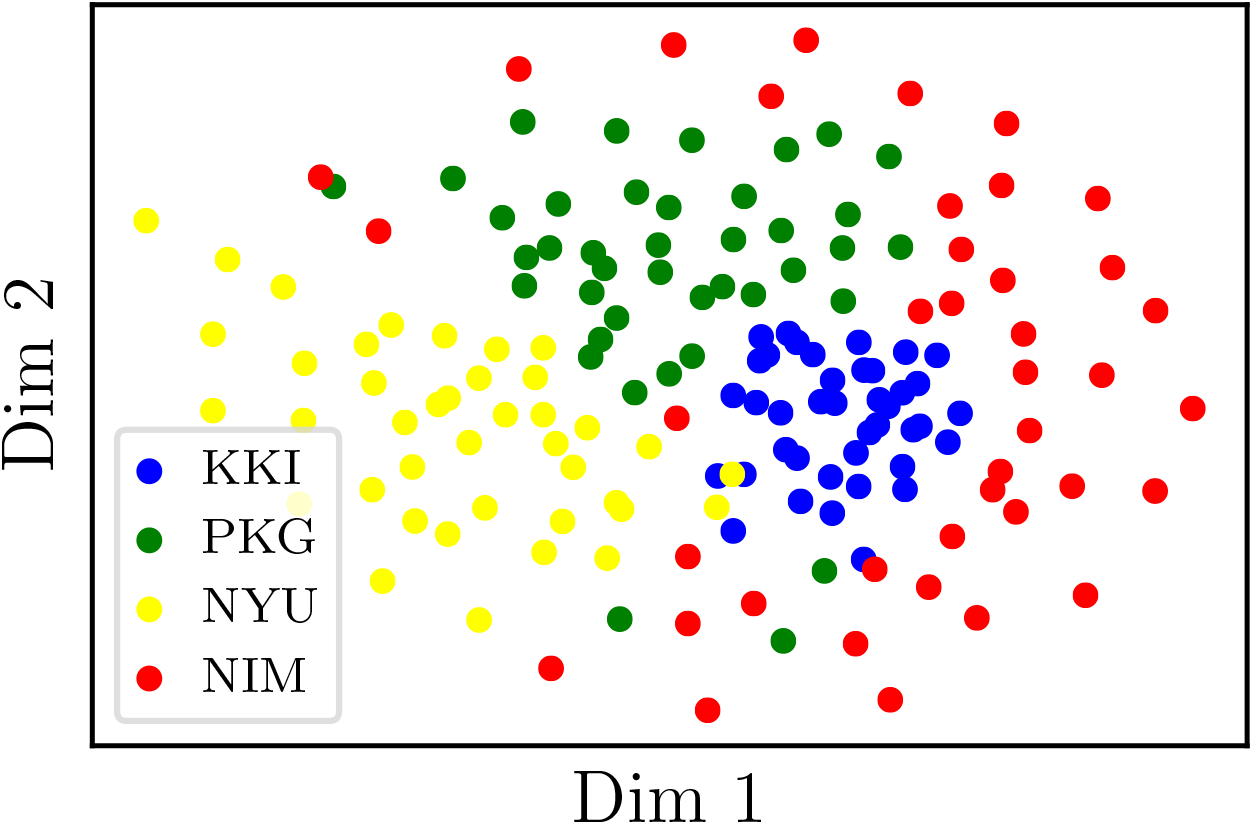
Two-dimensional embedding obtained after applying t-stochastic neighbour embedding to the logarithms of non-harmonized matrices. (KKI: Kennedy Krieger Institute, Johns Hopkins University; NYU: New York University Child Study Center; PKG: Peking University; NIM: NeuroIMAGE sample).

**Figure 4.**
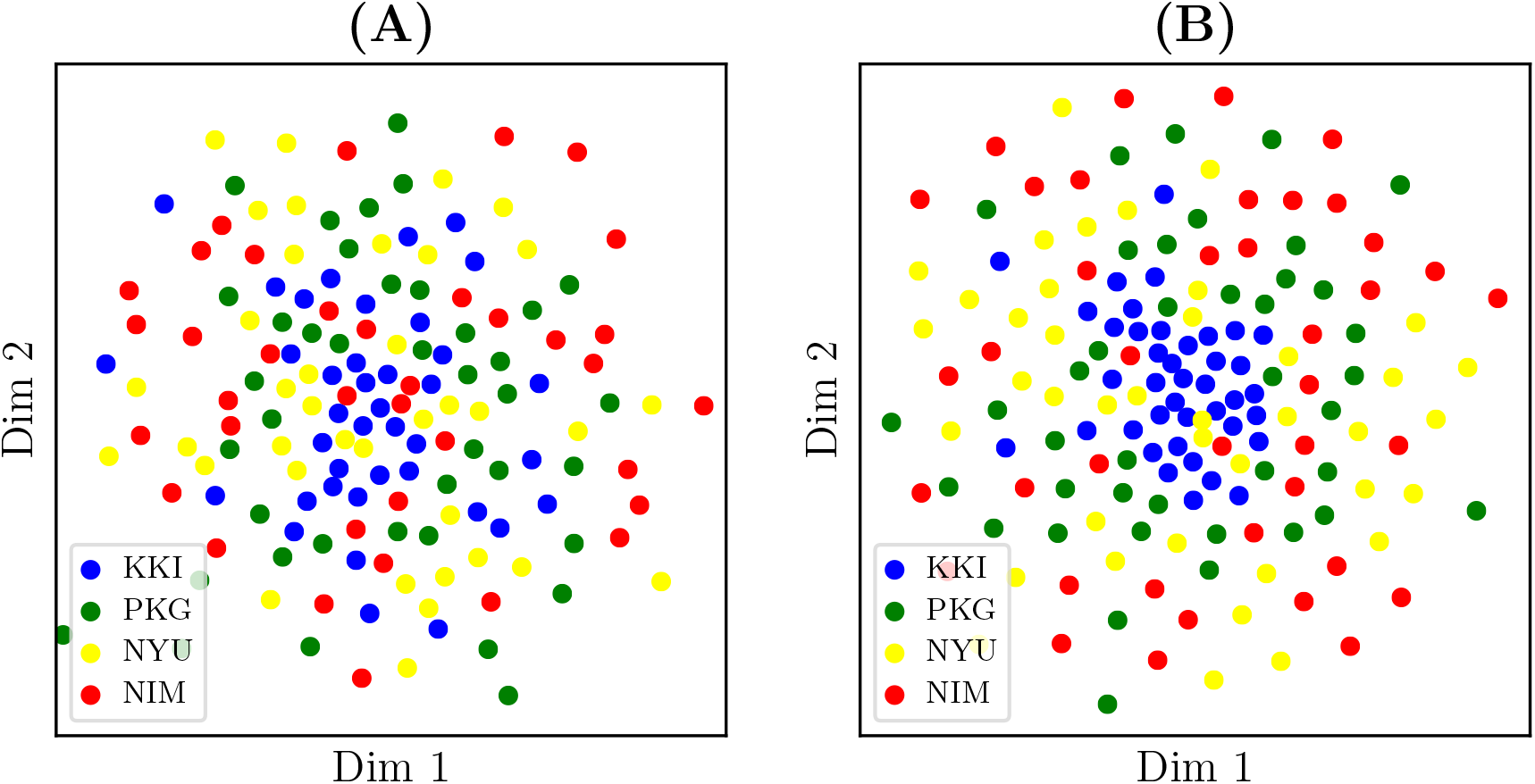
**(A)** Two-dimensional embedding obtained after applying t-stochastic neighbour embedding to the logarithms of matrices transformed under matrix whitening. **(B)** Two-dimensional embedding obtained after applying t-stochastic neighbour embedding to the logarithms of matrices transformed under Rigid Log-Euclidean Translation to the global mean 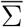. (KKI: Kennedy Krieger Institute, Johns Hopkins University; NYU: New York University Child Study Center; PKG: Peking University; NIM: NeuroIMAGE sample).

### 3.2 Numerical assessment of harmonization properties

Two harmonization properties were required from the beginning: preservation of intra-site geodesic distances, thus preserving biological variability (and some residual site bias), and the imposition of a common Fréchet mean. In the case of 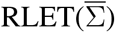, this Fréchet mean is the global mean 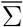, and in the case of RLET(*I*) and MW, the identity matrix *I*. To check that the first property is fulfilled, we have computed pair-wise distance difference matrices 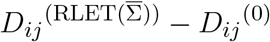 (with RLET(*I*) giving the same results by construction) and *D*_*ij*_^(MW)^ − *D*_*ij*_^(0)^ (in this latter case, using *D*_*ij*_^(PT)^ would give visually the same result, since *D*_*ij*_^(PT)^ ≈*D*_*ij*_^(MW)^, as proved before). We have found that diagonal blocks (differences in intra-site distances) vanish identically (Figure 5A and 5B). Regarding the second property, we have computed: on one side, pair-wise LERM distances between site means and between the global mean and these site means, before RLET(*I*) and 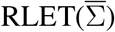, and after 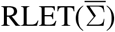 harmonization; and on the other side, pair-wise AIRM distances between site means and global mean before MW harmonization, pair-wise AIRM distances between site means and the identity matrix after MW harmonization, and pair-wise LERM distances between site means and the identity matrix after RLET(*I*) (Figures 5C, 5D, 5E, 5F). After harmonization these pair-wise distances vanish identically.

**Figure 5.**
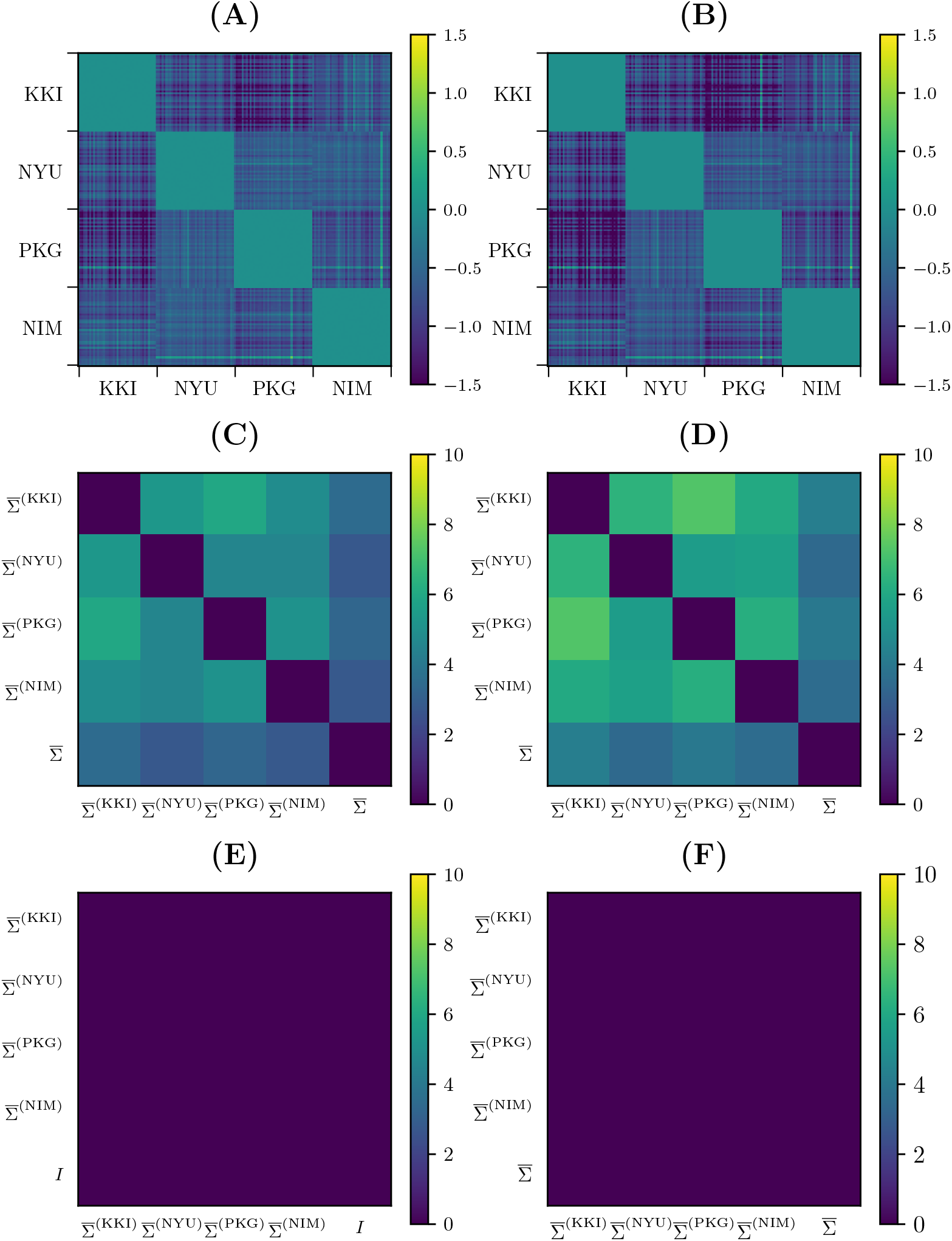
**(A)** Difference in pair-wise AIRM distances between original matrices and matrices transformed under matrix whitening. **(B)** Difference in pair-wise LERM distances between original matrices and matrices transformed under Rigid Log-Euclidean Translation (RLET). **(C)** Pair-wise AIRM distances between site Fréchet means 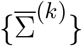 and the global mean 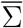. **(D)** Pair-wise LERM distances between site Fréchet means 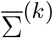 and the global mean 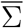. **(E)** Pair-wise AIRM distances between site Fréchet means computed after matrix whitening and the identity matrix *I*. **(F)** Pair-wise LERM distances between site Fréchet means computed after 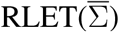 and the global mean 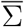. (KKI: Kennedy Krieger Institute, Johns Hopkins University; NYU: New York University Child Study Center; PKG: Peking University; NIM: NeuroIMAGE sample).

### 3.3 Entry-wise ADHD/HC differences

The process of performing 100 separate experiments, described in subsection 2.3, to detect entry-wise differences, has been applied, with the same randomly pooled subjects, to original, 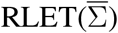-harmonized, RLET(*I*)-harmonized and MW-harmonized matrices (PT has been disregarded because of its approximate equivalence to MW). Therefore, we have obtained four frequency matrices *F*_*ij*_^(0)^, 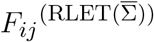, *F*_*ij*_^(RLET(*I*))^ and *F*_*ij*_^(MW)^, respectively, which are shown in Figure 6. Results on the sensitivity (defined also in subsection 2.3) we have obtained for *F*_*ij*_^(0)^, 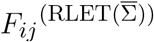,*F*_*ij*_^(RLET(*I*))^ and *F*_*ij*_^(MW)^ are shown in Table 1 between parentheses, together with the number of detected significant differences depending on their frequency of appearance in the frequency matrices.

**Table 1.** Number of detected anomalies using original matrices, harmonized matrices under Rigid Log-Euclidean Translation (RLET) to the global mean 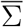, harmonized matrices under RLET to the identity matrix *I*, and harmonized matrices under matrix whitening, according to the chosen frequency threshold. Sensitivities are found between parentheses.

| Threshold | Original | RLET($\bar{\Sigma}$) | RLET($I$) | MW |
| --- | --- | --- | --- | --- |
| $F_{ij} \geq 1$ | 2019 | 2125 | 1753 | 2274 |
| $F_{ij} \geq 2$ | 188 (9.31%) | 213 (10.02%) | 154 (8.78%) | 262 (11.52%) |
| $F_{ij} \geq 3$ | 21 (1.04%) | 33 (1.55%) | 14 (0.80%) | 52 (2.3%) |
| $F_{ij} \geq 4$ | 2 (0.10%) | 8 (0.38%) | 1 (0.06%) | 9 (0.40%) |
| $F_{ij} \geq 5$ | 0 (0%) | 4 (0.19%) | 1 (0.06%) | 1 (0.04%) |

**Figure 6.**
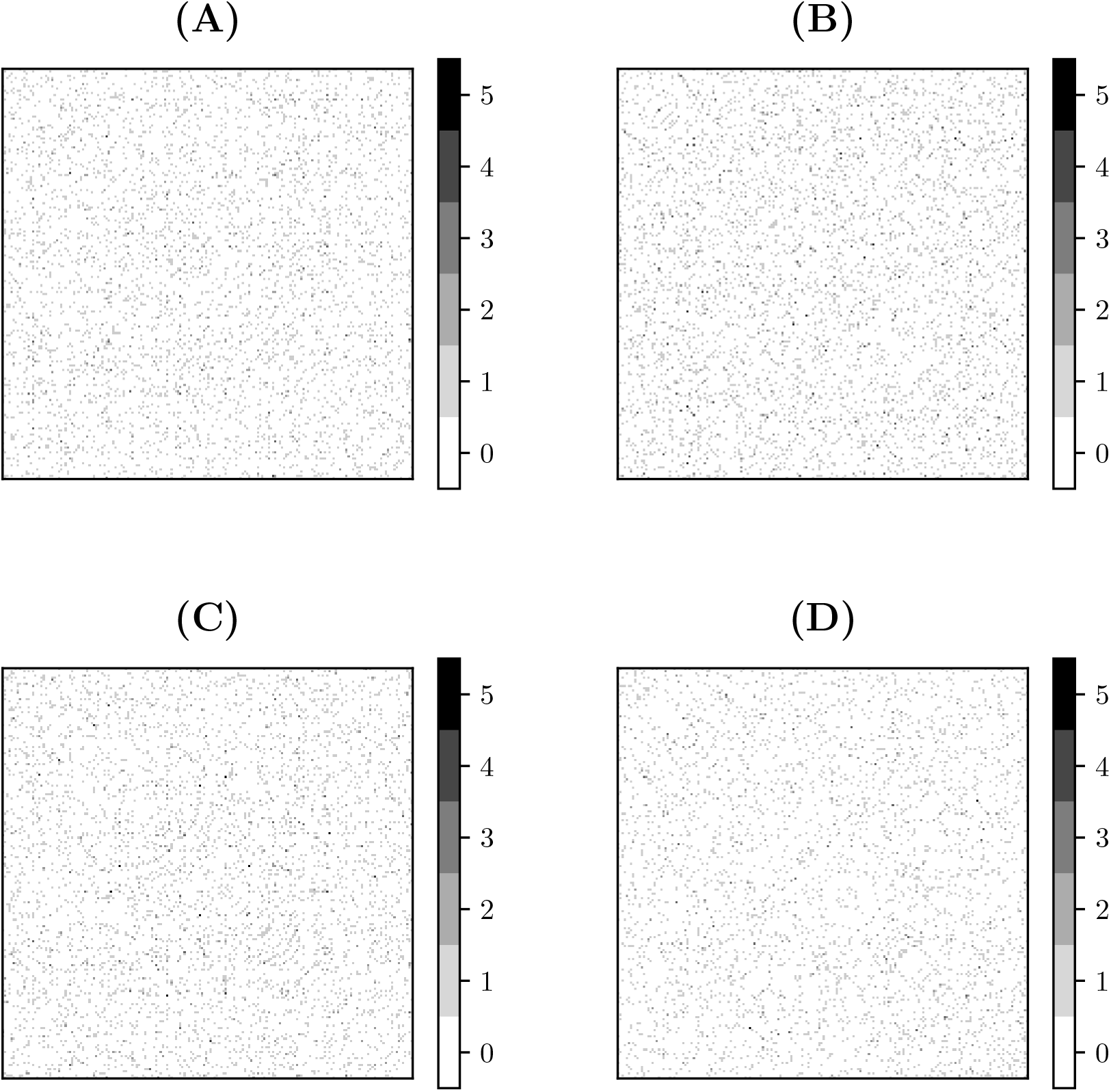
Frequency matrices *F*_*ij*_ obtained by performing 100 permutation experiments with the same randomly pooled subjects, binarizing each experiment’s p-value matrix at *p <* 0.001, and summing these resulting matrices for: **(A)** original functional connectivity (FC) matrices; **(B)** FC matrices transformed under matrix whitening; **(C)** FC matrices transformed under Rigid Log-Euclidean Translation (RLET) to the global mean 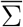; **(D)** FC matrices transformed under RLET to the identity matrix.

### 3.4 Intra-network and inter-network anomalies distribution

In the following, we will refer to significant differences in connection strength between ADHD and HC subjects as ‘anomalies’. To have a global view of the distribution of anomalies in terms of inter- and intra-network interaction, we have constructed a 2-dimensional plot that reflects the relative involvement of the different resting-state networks in these interactions.

This 2-dimensional plot is a 10×10 matrix (one row and one column per functional component) constructed as follows: first, we have filled the matrix by assigning to each entry *ij* the number of anomalies connecting functional component *i* with functional component *j* (therefore, diagonal elements represent intra-network anomalies), with the matrix being symmetric since we do not have a sense of directed connection; afterwards, we have normalized each row (say, for example, DMN), excluding diagonal elements, by using the total number of inter-network anomalies that involve the component represented by that row (DMN); finally, in the last step, we have normalized the diagonal by dividing each diagonal entry by the sum of all diagonal entries (that is, the total number of intra-network anomalies). At this point, we are left with a matrix where each row, excluding diagonal entries, adds up to 1, and in turn, diagonal entries add up to 1. One has to interpret this matrix in the following manner: when taking, for example, the row corresponding to DMN, the entry DMN-FP is the percentage of inter-network anomalies involving DMN that end up connected to FP; when considering DMN-DMN, it is the percentage of intra-network anomalies that are found within DMN. Therefore, we have summarized in a plot the relative contribution of every functional component to the inter-network anomalies involving another specific functional component. One has to bear in mind that DMN-FP (row-column) does not have the same interpretation as FP-DMN and, indeed, their values are different in general (after normalizing per row, the matrix is no longer symmetric): the number of anomalous connections between DMN and FP is fixed, but their relative contribution to the total number of inter-network anomalies involving DMN and to the total number of inter-network anomalies involving FP need not be the same.

We have plotted these matrices for the anomalies obtained using the original, the 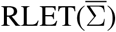 -harmonized, the RLET(*I*)-harmonized and the MW-harmonized functional connectomes (Figure 7). They have been computed with the anomalous connections identified by thresholding their corresponding frequency matrices at *F*_*ij*_ ≥ 2 to have a large enough number of samples (Table 1). Furthermore, we have computed differences between the plots corresponding to original and 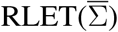, between RLET(*I*) and MW, and between 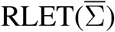 and RLET(*I*); afterwards, we have binarized the resulting difference plots at an absolute entry-wise difference ≥ 15%, thus allowing the detection of important changes in the distributions and assessing harmonization and manifold location effects. These results are shown in Figure 8.

**Figure 7.**
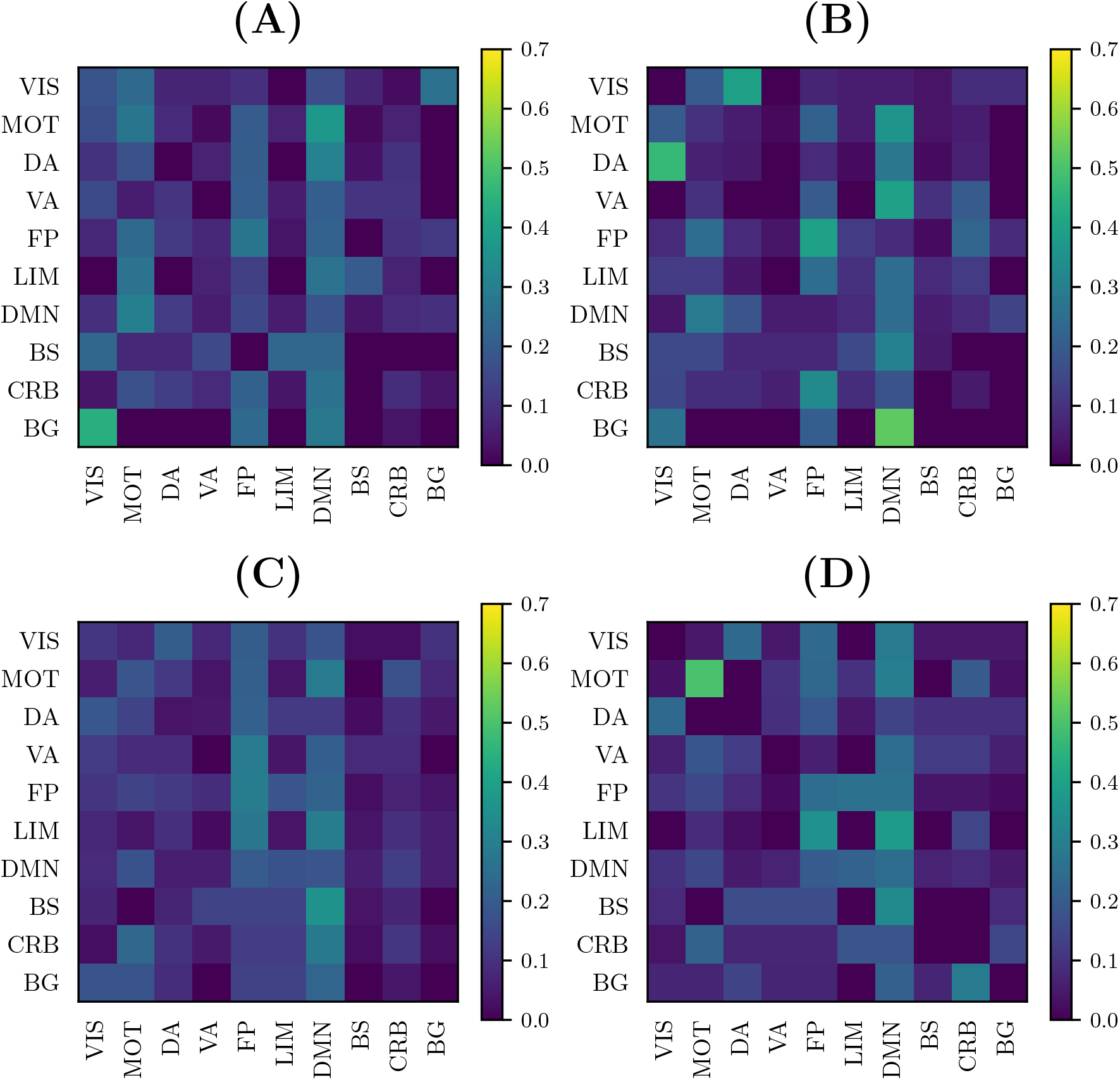
Anomaly distribution plots according to the involved functional components obtained as described in subsection 3.4 using: **(A)** original matrices; **(B)** matrices transformed under Rigid Log-Euclidean Translation (RLET) to the global mean 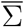; **(C)** matrices transformed under matrix whitening; matrices transformed under RLET to the identity matrix. (VIS: visual network; MOT: sensorimotor network; DA: dorsal attention network; VA: ventral attention network; FP: frontoparietal network; LIM: limbic network; DMN: default mode network; BS: brain stem; CRB: cerebellum; BG: basal ganglia).

**Figure 8.**
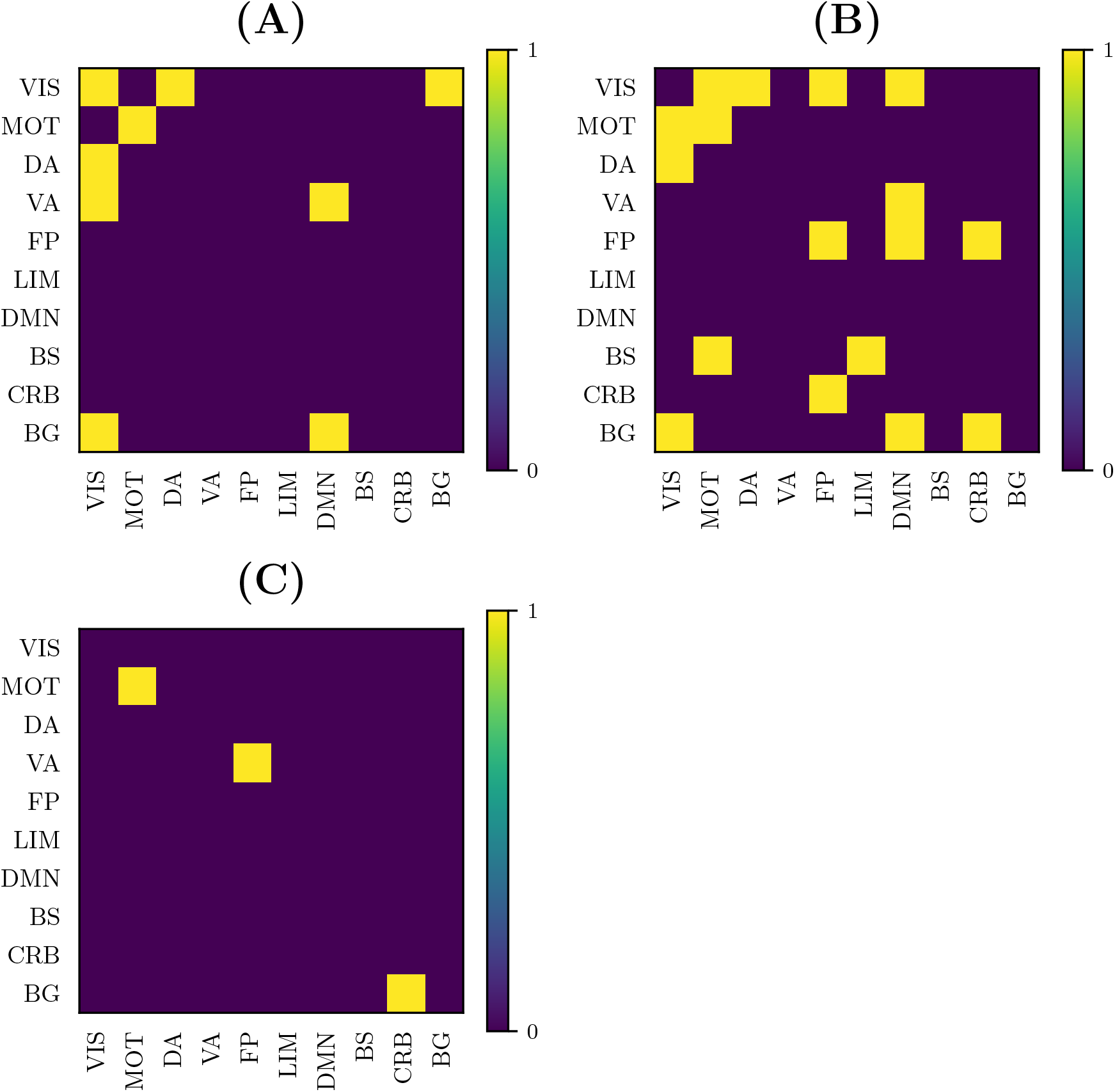
Binarized entry-wise absolute differences greater than 15% between plots in Figure 6 for: **(A)** original (non-harmonized) matrices and matrices harmonized under Rigid Log-Euclidean Translation (RLET) to the global mean 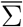; **(B)** RLET to 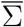 and RLET to the identity matrix *I* harmonization approaches; **(C)** matrix whitening and RLET to *I* harmonized matrices. (VIS: visual network; MOT: sensorimotor network; DA: dorsal attention network; VA: ventral attention network; FP: frontoparietal network; LIM: limbic network; DMN: default mode network; BS: brain stem; CRB: cerebellum; BG: basal ganglia).

Even though a more involved interpretation of these results will be made in the Discussion section, we highlight that discrepancies between 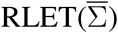 and RLET(*I*) signal distortions induced by relocating matrices to the neighbourhood of *I*, since relative positions between matrices in Log-Euclidean space are exactly the same by construction. Therefore, in the following, we will focus on the comparison between non-harmonized and 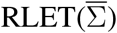-harmonized matrices.

### 3.5 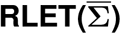-harmonization results

At this point, we focus on results at a frequency *F*_*ij*_ ≥ 3 to allow a more precise comparison between the results obtained from non-harmonized and 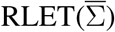-harmonized matrices. In this case, we have obtained 21 significant inter-network connections using non-harmonized matrices; and 32 inter-network and 1 intra-network anomalies after transforming original matrices under the 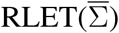 framework. We have represented the resulting anomalous connectomes in Figures 9,10.

**Figure 9.**
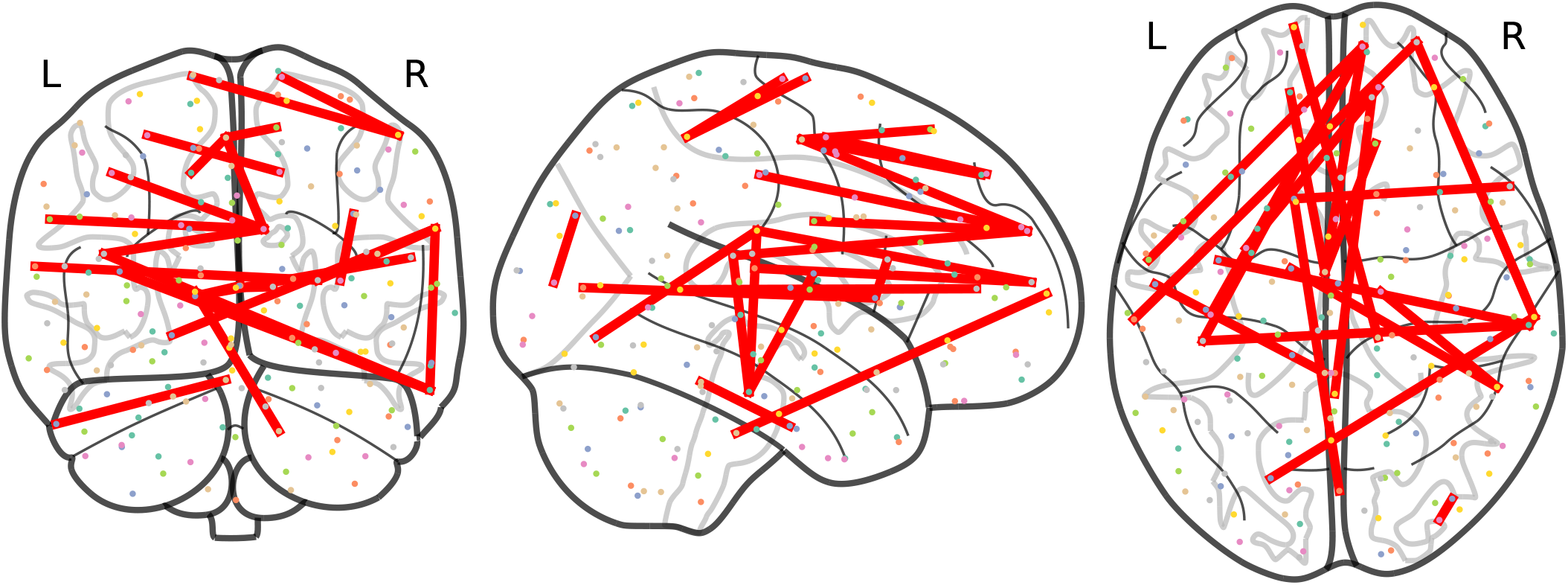
Anomalous connections obtained by thresholding the frequency matrix for non-harmonized matrices at *F*_*ij*_ ≥ 3. In total, the 21 anomalous connections are displayed. Notice the lack of anomalies involving the posterior parietal cortex and the cerebellum.

**Figure 10.**
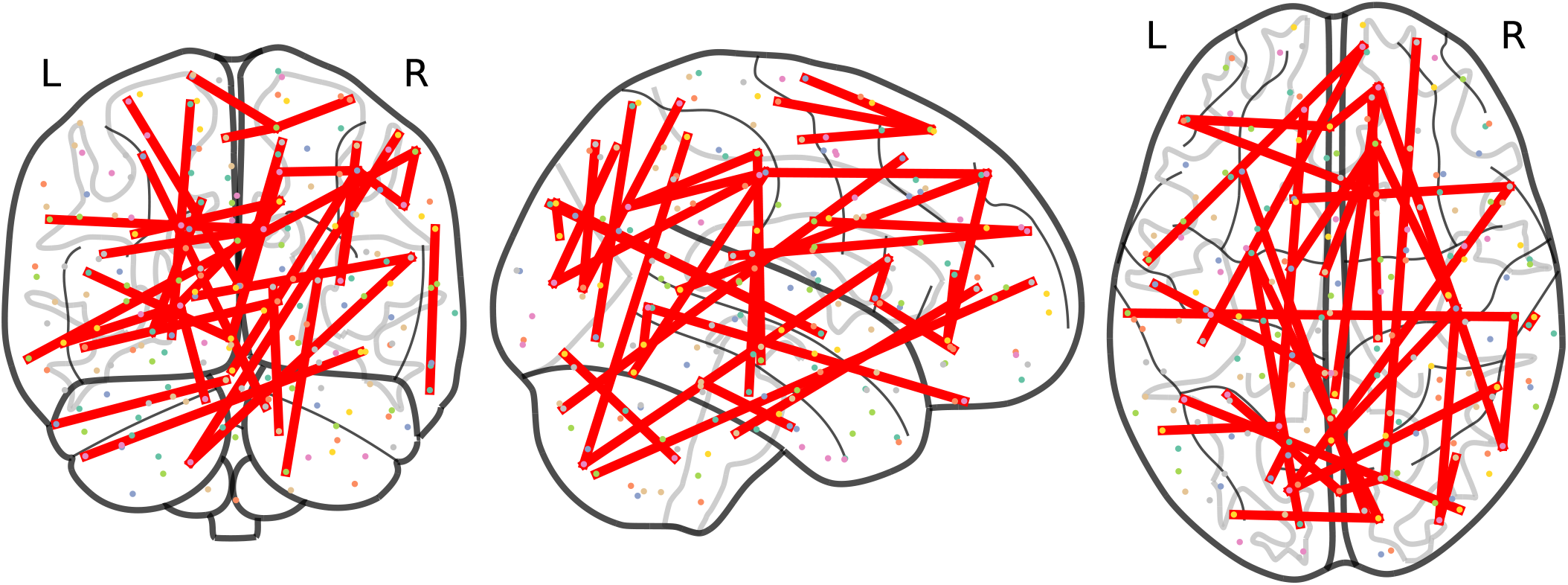
Anomalous connections obtained by thresholding the frequency matrix for matrices harmonized under Rigid Log-Euclidean Translation to the global mean 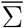 at *F*_*ij*_ ≥ 3. The 33 detected anomalies are displayed.

Moreover, we have plotted two histograms, one for non-harmonized and the other for 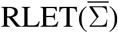-harmonized matrices, displaying how the distribution of abnormal connections involving particular functional components varies according to the frequency threshold (Figure 11). When looking at the histograms, one can see that the distribution of anomalies according to the involved functional component is better preserved when changing the considered frequency level in the case of 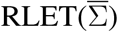 than in the non-harmonization approach.

**Figure 11.**
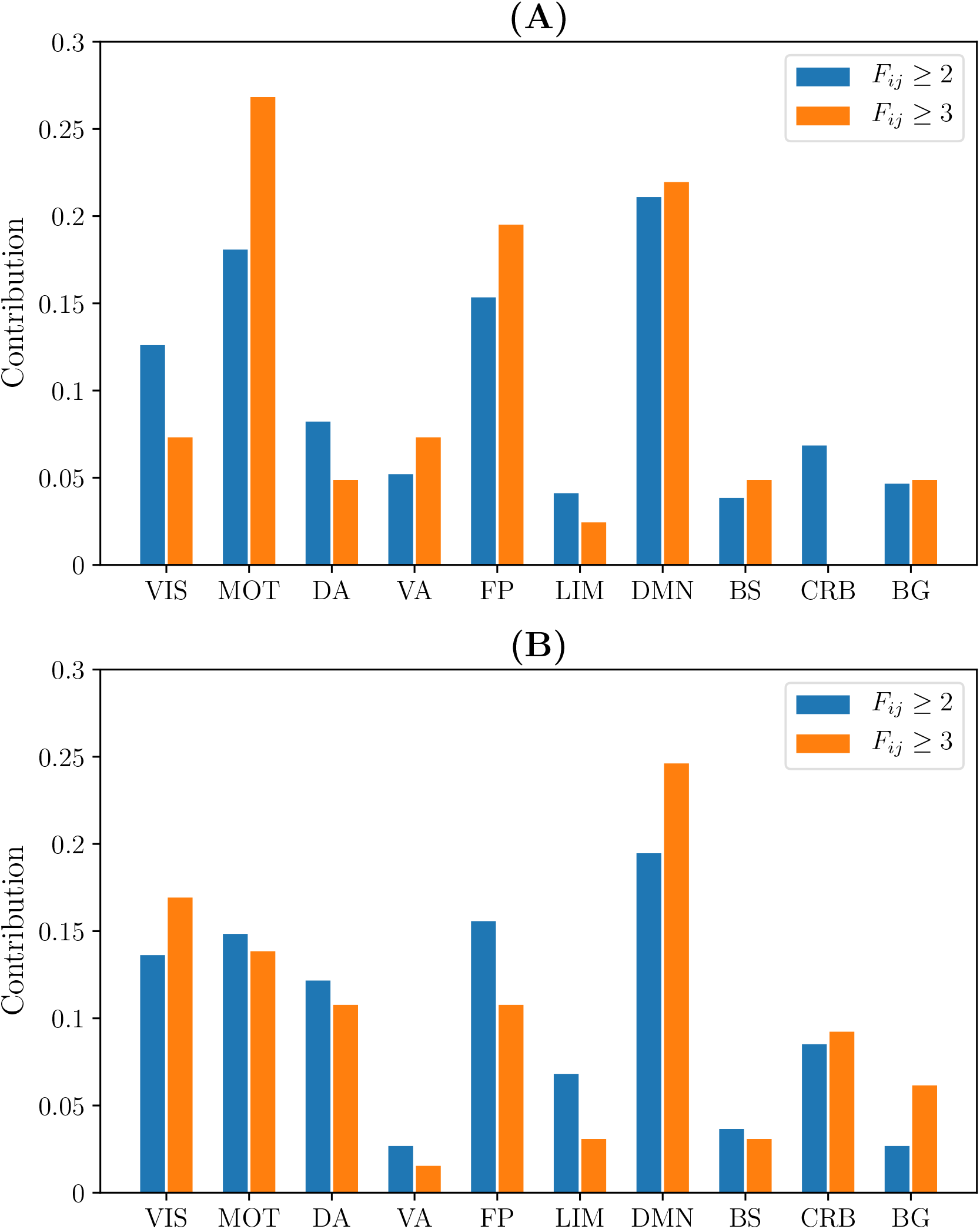
Relative contribution of functional components to the total number of detected anomalies by thresholding frequency matrices at *F*_*ij*_ ≥ 2 and *F*_*ij*_ ≥ 3 corresponding to: **(A)** the non-harmonization approach; **(B)** harmonization under Rigid Log-Euclidean Translation to the global mean 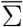. (VIS: visual network; MOT: sensorimotor network; DA: dorsal attention network; VA: ventral attention network; FP: frontoparietal network; LIM: limbic network; DMN: default mode network; BS: brain stem; CRB: cerebellum; BG: basal ganglia).

## 4 DISCUSSION

### 4.1 Imaging-site bias

Results from pair-wise distances computation (Figure 1) and low-dimensional embedding (Figures 3 and 4) of functional connectivity matrices show that there is an imaging-site-biased distribution of these matrices in the manifold. In particular, one can distinguish between an intra-site bias effect and an inter-site bias effect. Intra-site bias is clearly visible when looking at diagonal blocks of pair-wise distances plots: intra-site distances are significantly different between KKI and NIM matrices (interestingly, the NIM sample comes in its turn from 3 different imaging sites (ADHD-200 Consortium, 2017), what could explain its large intra-site variability). Instead, one would expect the intrinsic biological variability to be approximately the same across sites when the number of subjects is large enough. Inter-site bias is reflected in the clustered distribution of matrices in the manifold that makes possible to clearly identify visually the imaging site in the tSNE low-dimensional embedding (recall that this low-dimensional embedding is, with high probability, displaying LERM distance similarities). When harmonizing using the three different methods, inter-site distances were generally reduced (Figure 1), which is the expected effect of removing site-clustering behaviour.

The clustered distribution in the manifold (Figure 3) points directly to the fact that entry-wise statistical comparisons could not be completely reliable unless the site effect is previously subtracted (Figure 4), since what are usually targeted in these statistical tests are entry-wise differences. Because of the nature of the permutation testing approach targeting these differences, large entry-wise differences coming from the original clustered and distant distribution of the matrices might be masking true and more subtle biological differences, or even giving rise to wrong salient entries. Geometry-grounded frameworks transform the matrices’ entries in such a way that they can still be regarded as covariances (resulting matrices belong to the SPD manifold, as opposed to ComBat outputs, for example), that removes (partially) multi-site effects on entry-wise differences, and therefore that allows a more unbiased study of significance of these individual differences or anomalies.

Regarding the detection of significant differences, in terms of sensitivity RLET(*I*) is the harmonization method that performs the worst (Table 1) at all frequency thresholds. Nevertheless, only by using harmonization approaches have we reached the *F*_*ij*_ = 5 level. On the other hand, site means transform as expected, becoming the identity matrix *I* for RLET(*I*) and MW, and the global mean 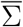 in the case of 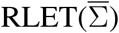, since pair-wise distances vanish. Intra-site distances are also preserved as intended but, as a consequence, intra-site bias is also retained. However, as a first approximation to imaging-site harmonization, our aim was to operate in the least distortive manner. One cannot know to which extent true biological variability would be artificially altered by modifying intra-site distances, even though it is clear that there is an imaging-site contribution to the observed dispersion. Notice however that when looking at Figure 4A, it seems that MW has been able to correct differences in dispersions, as opposed to RLET (Figure 4B). This impression is an artifact coming from the fact that the low-dimensional embedding has been obtained by using LERM distances, both for MW and for RLET. MW is based on the AIRM framework and, therefore, the low-dimensional embedding, using LERM distances as the metric, has not been able to capture the expected intra-site AIRM distance preservation behavior. In fact, if one considers Figure 1, one can see that different dispersions around the site mean have not been corrected for MW (Figure 1C), since diagonal blocks exhibit different average intra-site distances in the same way as RLET-transformed ones (Figure 1D).

Figure 8 provides evidence favouring harmonization, but also evidence for rejecting harmonization at the identity matrix. Testing for the difference between these distribution matrices and requiring entry-wise coincidence within ±15% is strict, bearing in mind that we are dealing with relative contributions of connections among and within 10 different components, and the number of samples is considerably small. Therefore, at this point, looking at the differences between RLET(*I*) and MW (only 3/100 entries display absolute differences larger than 15%), we can claim that harmonization is indeed working in its task of homogenizing results. However, taking into account that the only difference between RLET(*I*) and 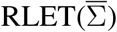 is the location of the Fréchet mean (relative positions between functional connectivity matrices in the Log-Euclidean space are exactly preserved by construction), and considering the dissimilarities between distributions, we can affirm that: location in the manifold is important (as we had presumed in the Methods section, and motivating our construction of 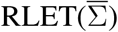 and that transporting matrices to the identity matrix is too distortive in our setting. As a consequence, and given that discrepancies in anomaly distributions between original and 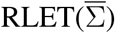 are three-fold (9/100) the ones observed by harmonizing with two different approaches at identity (3/100), we have focused our attention in the following subsection on the effect of 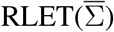-harmonization on the original functional connectivity matrices, since these (possibly biologically relevant) discrepancies are, at least partially, due to harmonization.

Finally, given the results pointing to a considerably large intra-site bias, we can propose a modification to 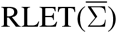:

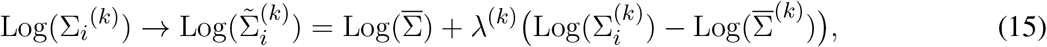

consisting on the introduction of a site-dependent rescaling parameter *λ*^(*k*)^ *>* 0, which has the effect of rescaling intra-site distances by a factor of *λ*^(*k*)^ while still enforcing all site Fréchet means to be the global mean 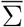. Tuning *λ*^(*k*)^ would allow, for example, to empirically adjust mean intra-site distances (and, thus, intra-site variability), so that different sites display approximately the same dispersion around 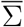. A first approximation choice for *λ*^(*k*)^ could be therefore:

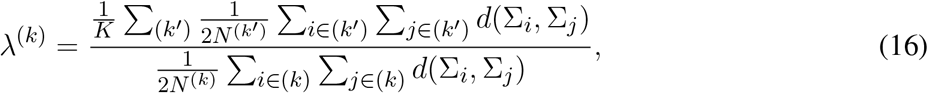

that is, the ratio between the mean of all intra-site pair-wise distances (regardless of *k*) and the mean of intra-site-*k* distances.

### 4.2 Abnormal functional connectivity findings

#### 4.2.1 Large-scale networks in ADHD

By thresholding frequency matrices at *F*_*ij*_ ≥ 2 one can obtain a large enough number of anomalies to construct the distributions shown in Figure 7. This approach enables to obtain a coarse-grained picture of anomalous interactions and, therefore, to take into account anomalous engagement between large-scale resting-state networks.

Although there is a large amount of pair-wise interactions, the most descriptive findings are obtained by considering relevant functional components that do not have anomalous inter- or intra-network interactions. Using non-harmonized matrices, we have found that at *F*_*ij*_ ≥ 2 there is an absence of intra-DA and intra-VA anomalous connections. In the case of 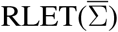 -harmonized matrices, we see an absence of intra-VIS and intra-VA anomalies, and an absence of DA-VA and VIS-VA inter-network abnormal interaction. In both cases, BG engage abnormally selectively with DMN, FP and VIS networks. In resting-state functional connectivity, in general, findings do not support an involvement of VA in ADHD, even though authors do not find this evidence to be conclusive (Castellanos and Proal, 2012). On the contrary, DA involvement in ADHD is well-established (Dickstein et al., 2006; Rubia, 2011). Absence of intra-DA alterations under the non-harmonization framework could be therefore interpreted as a deficit of the framework. Intra-VIS alterations, both hyper-and hypoactivations, have been found in task-based settings, depending on the task (Dillo et al., 2010; Schneider et al., 2010; Vance et al., 2007). One study found altered intra-VIS resting-state functional connectivity by using ADHD-200 resting-state data (Kessler et al., 2014) (correlating with results from the analysis of non-harmonized matrices, where intra-VIS contributions are found). Authors highlight the importance of examining carefully the impact of the visual component in ADHD and, in particular, its relation to attention (Castellanos and Proal, 2012). To our opinion, given the importance (in terms of contribution to anomalies) that 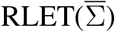-harmonization gives to VIS, the absence of intra-network findings is descriptive, in the sense that it is difficult to understand that the algorithm did not discover intra-network anomalies because of sensitivity regarding the visual component.

#### 4.2.2 Pathophysiology of ADHD

Although there is a growing consensus that large-scale brain networks are involved in ADHD, the number of anomalous connections that remain as significant when thresholding their frequency at *F*_*ij*_ ≥ 3 does not allow an interpretation in terms of network interactions, but rather an interpretation in terms of pathways, (parts of) circuits or pair-wise relevant connections. This interpretation makes possible to probe the fine-grained pathophysiology of ADHD from a functional point of view, and to correlate our findings with established results.

##### 4.2.2.1 Prefrontal cortex

It has been hypothesized for a long time that ADHD is a disorder of the prefrontal cortex (PFC). The motivation underlying the hypothesis of involvement of PFC is that one of its most important functions is behavioural control, the principal impairment presented by ADHD patients. The circuitry that has been found to be critical in the neurobiology of ADHD are the cortico-striatal-thalamo-cortical (CSTC) loops. These closed loops transmit cortical inputs to the thalamus via the striatum, to be reprojected back to the cortical region. Two of these loops play a key role in ADHD: the dorsolateral prefrontal circuit, with the dorsolateral prefrontal cortex (DLPFC) and the (dorsolateral) caudate nucleus forming the cortico-striatal pathway, and the orbitofrontal circuit, composed of the orbitofrontal cortex (OFC) and the (ventromedial) caudate nucleus pathway. The dorsolateral prefrontal circuit is a central element in cognitive control, and the orbitofrontal circuit is heavily implicated in reward processing, whose impairment is thought to provoke impulsive behaviour in ADHD. On top of these ones, another component that has been added more recently to ADHD PFC anomalous connections is the fronto-cerebellar circuitry. The cerebellum is thought to be involved in diverse functional impairments of ADHD patients, such as working memory, attention, or the construction of temporal expectations. For a detailed description of the involvement of these circuits in ADHD, we refer the reader to (Durston et al., 2011) and references therein.

Remarkably, the analysis of 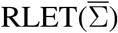-harmonized matrices points directly to these circuits: we have found anomalous connectivity between the right caudate nucleus and the left OFC; between right cerebellum (VIIb) and the right frontal pole; together with a connected chain comprising the left caudate nucleus, the right inferior frontal gyrus pars opercularis, left cerebellum (VIIb) and the right angular gyrus, with both right inferior frontal gyrus pars opercularis and right angular gyrus lying on DLPFC. Therefore, we have found evidence for: 1) anomalies in frontostriatal paths, both in the OFC and DLPFC circuits; 2) an abnormal connectivity in a fronto-cerebellar path (right cerebellum - right frontal pole); 3) a simultaneous anomalous engagement of fronto-cerebellar and frontostriatal circuits with the same cortical region, DLPFC (left cerebellum - right angular gyrus, right inferior frontal gyrus pars opercularis - left caudate nucleus); and 4) more specifically, a simultaneous anomalous engagement of a fronto-cerebellar circuit and a frontostriatal circuit involving exactly the same cortical node (right inferior frontal gyrus pars opercularis, DLPFC), and, thus, a precise anomalous overlap of different circuitry components. Results of non-harmonized matrices also feature the anomalous connection between the left caudate nucleus and the right inferior frontal gyrus pars opercularis. In this case, we would only have evidence for the involvement of the DLPFC circuit. Furthermore, it is important to mention that non-harmonized matrices have not signaled any anomaly involving cerebellar nodes, which is undesirable given the pervasive evidence of their implication in ADHD. In contrast, 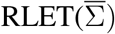-harmonization provides evidence of abnormal functional connectivity of the cerebellum with somatomotor, default mode network and visual nodes, apart from the PFC ones, pointing to the multi-dimensional association to ADHD pathophysiology.

##### 4.2.2.2 Visual nodes

It has been established that ADHD patients have worse performances when tested for visual processing speed and visual short-term memory in comparison to matched healthy controls (Low et al., 2018). These abnormal performances signal a perceptual deficit and, more concretely, impairments in early visual information processing (Alqahtani et al., 2019; Papp et al., 2020).

One of the most remarkable differences between the anomalies spotted using the original matrices and the 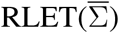-harmonized ones is the importance that the analysis of the latter matrices gives to visual network nodes, in terms of contribution to the total number of detected anomalies. Specifically, there is an interesting and insisting presence of anomalous connections between visual network nodes and nodes found in the posterior parietal cortex (PPC), belonging to the dorsal attention network (DA). Precisely, these connections fill the void found in Figure 9. It has been found that the interaction between the occipital cortex and DA is involved in maintaining attention and suppressing attention to irrelevant stimuli (Shulman et al., 2009; Capotosto et al., 2009). Our findings signal specifically the anomalous connection of nodes lying on the most posterior part of DA and nodes of the visual cortex. One important pathway for visual information processing connects these brain regions: the dorsal stream. The dorsal stream and the ventral stream were proposed as two routes with differentiated functions in early processing of visual information (Goodale and Milner, 1992): the dorsal stream, running from the visual cortex to PPC, is typically thought to be involved in the construction of a detailed map of the visual field, spatial awareness, the detection and analysis of motion and the guidance of actions and coordination in space; on the other hand, the ventral stream, originating in the visual cortex and leading to the temporal lobe, is proposed to be related to visual identification and recognition.

The dorsal stream, which is tentatively being featured in our findings, has been demonstrated with clinical and experimental evidence to play an essential role in the ability of shifting spatial attention, that is, to disengage attention from a location and to engage attention to another location (Sciberras-Lim and Lambert, 2017). Although in ADHD patients there is a disregulation in attentional resources allocation, it is still not clear which dimensions of attention are affected by the disorder. The affectation of visuospatial attention is a controversial topic (Roberts et al., 2017), with findings pointing to opposite ways. Many authors point to the fact that given the wide spectrum of impairments that patients exhibit, their phenotyping by a particular dimension of attention might not be possible (Roberts et al., 2017). Nevertheless, what is widely accepted is that ADHD patients display a deficit in visuospatial working memory (van Ewijk et al., 2013), the capacity to maintain a representation of visuospatial information for a brief period of time (Vecera and Rizzo, 2003). In particular, visuospatial working memory is involved, for example, in retrieval and manipulation of recent images for orientation in space and for keeping track of moving objects. As we have explained, the dorsal stream plays a critical role in the integration of visual information to produce the egocentric spatial map on which the subject relies to detect motion or to guide his/her actions in space. Therefore, there is coherence between the impairment of visuospatial working memory in ADHD and the tentative appearance of the dorsal stream in our findings.

##### 4.2.2.3 Other relevant anomalies

Given the coherence between the results obtained after 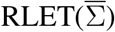-harmonization and established ADHD pathophysiology, we have reasons to think that default mode network (DMN), sensorimotor (MOT), visual (VIS), dorsal attention (DA) and frontoparietal (FP) nodes play very important roles in the disorder, and that the right lateral occipital cortex is a region that needs to be examined more carefully in subsequent analyses. When thresholding the frequency matrix at *F*_*ij*_ ≥4, we are left with 8 anomalous connections. Within these 8 anomalies, we can find the aforementioned OFC-striatum and fronto-cerebellar pathways, and 5 of them involve DMN. If we take the maximum threshold *F*_*ij*_ = 5, 4 of these previous 8 anomalies remain: left superior frontal gyrus (MOT) with right superior frontal gyrus (DMN), right postcentral gyrus (MOT) with right lateral occipital cortex superior division (DMN), right occipital pole (VIS) with right lateral occipital cortex superior division (FP), and right lateral occipital cortex superior division (DMN) with right supramarginal gyrus anterior division (DA). Interestingly, 3 out of 4 feature the same brain region, the right lateral occipital cortex superior division, being associated to DMN and FP functional networks, depending on the particular node of the region.

### 4.3 Conclusions

In conclusion, we have collected evidence that supports further research on multi-site dataset harmonization when using functional connectivity matrices. Specifically, we have shown that the distribution of matrices in the symmetric positive-definite matrices manifold is site-biased. Moreover, we have been able to discard two adapted domain adaptation approaches and to prove that our proposal 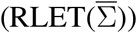 is the most reliable one in a clinical setting. Remarkably, functional analyses of ADHD-200 data after 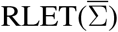-harmonization present a high correlation with neurobiological findings in ADHD. However, intra-site bias was not removed and, to our point of view, this issue needs to be addressed, possibly by using our modification of RLET as a first approximation. It would also be valuable to use a more extended dataset for further research, and to rely on more permutations in each permutation test to increase statistical power. Nevertheless, given our results, we think there are reasons for reanalyzing functional data from multi-site datasets corresponding to other disorders, given their pervasive use in functional neuroscience.

## CONFLICT OF INTEREST STATEMENT

The authors declare that the research was conducted in the absence of any commercial or financial relationships that could be construed as a potential conflict of interest.

## AUTHOR CONTRIBUTIONS

GS performed the study conception and design, as well as data analysis, code implementation and material and figures preparation. The first draft of the manuscript was written by GS. DP, GP and OC provided methodological guidance according to their role of master thesis’ supervisors and senior authorship, and revised the manuscript.

## FUNDING

This research has not been supported by any kind of funding.

## ACKNOWLEDGMENTS

We thank the reviewers for their useful and constructive comments and suggestions. We also thank the publisher Frontiers for the partial publication fee waiver.

## DATA AVAILABILITY STATEMENT

The code and the functional connectivity matrices used in this work, along with the p-value and frequency matrices generated after statistical experiments are available at Simeon (2021). The conversion from ROI number to anatomical location can be found along with the CC200 parcellation at Craddock (2011).

## ANNEX

### Equivalence between parallel transport and matrix whitening

In general, matrix whitening and parallel transport using 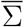 as the reference point are different transformations. However, we show here that under the condition 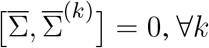, both frameworks are equivalent. To this aim, consider the following property regarding the unique positive-definite square root of the product of two *n* × *n* SPD matrices:

#### Property

Let *A* and *B* be two *n* × *n* SPD matrices that commute, i.e., [*A, B*] ≡ *AB* − *BA* = 0. Then 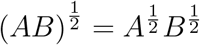.

#### Proof

Considering that the unique SPD square root of some SPD matrix *X* can be computed as 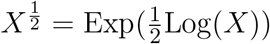, and taking into account that, given two SPD matrices *A* and *B* such that [*A, B*] = 0, Exp(*A* + *B*) = Exp(*A*) Exp(*B*) and Log(*AB*) = Log(*A*) + Log(*B*):

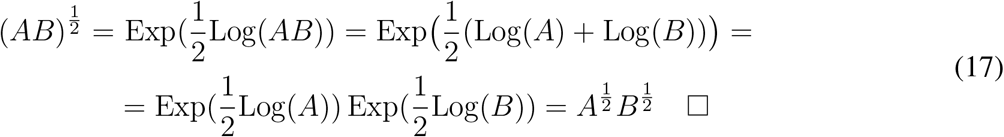

Now, recall that the overall parallel transport transformation (including the last matrix whitening step) when using as the reference point Σ_0_ the global mean 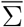 (Yair et al., 2019) is (subsection 2.4.2):

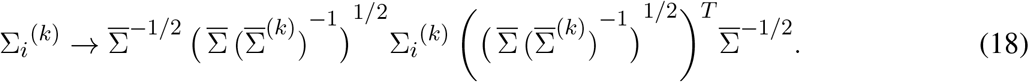

Supposing that 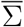 and 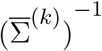 commute, by virtue of the previously stated property and taking into account that given two matrices *A* and *B*, (*AB*)^*T*^ = *B*^*T*^ *A*^*T*^ :

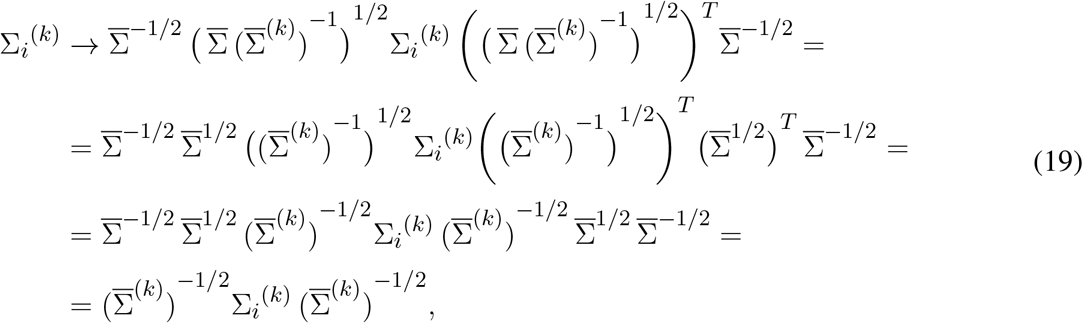

where in the second step we have used the facts that for a SPD matrix *A*, we take *A*^1*/*2^ to be also SPD and thus satisfies (*A*^1*/*2^)^*T*^, along with ((*A*)^−1^)^1/2^ = (*A*) ^−1^, with (*A*)^−1*/*2^ also SPD, and then, ((*A*)^−1/2^)^*T*^ = (*A*) ^−1/2^. One can conclude that under vanishing commutators 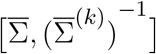, the transformation reduces to matrix whitening. Since 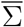 and 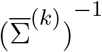 belong to *GL*(*n*), the requirement 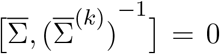 is equivalent to 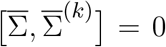. Therefore, when site means 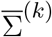 commute with the global mean 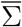, parallel transport and matrix whitening represent equivalent frameworks.

### Properties of Rigid Log-Euclidean Translation

As mentioned in the description of Rigid Log-Euclidean Translation in subsection 2.4.3, the transformation preserves intra-site geodesic distances under the LERM framework and displaces the matrices in such a way that their transformed site mean 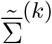 is the global mean 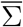, for all sites *k*.

By using the definition of LERM geodesic distance (7) for matrices belonging to the same site *k* and modified according to 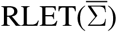 transformation (14) one finds

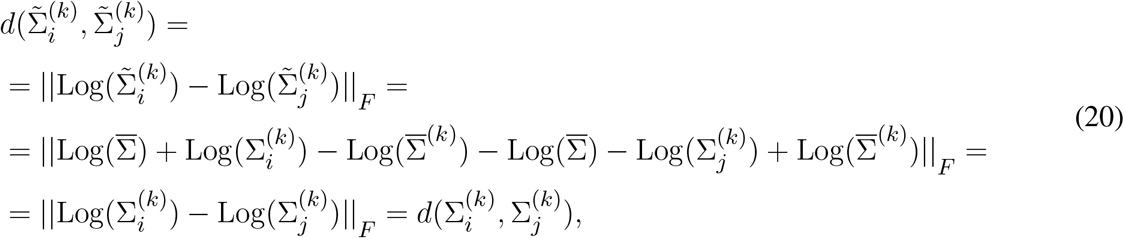

meaning that intra-site geodesic distances are preserved as expected.

It is also straightforward to prove that 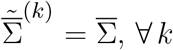. Considering the definition of site mean (12) for matrices transformed according to (14):

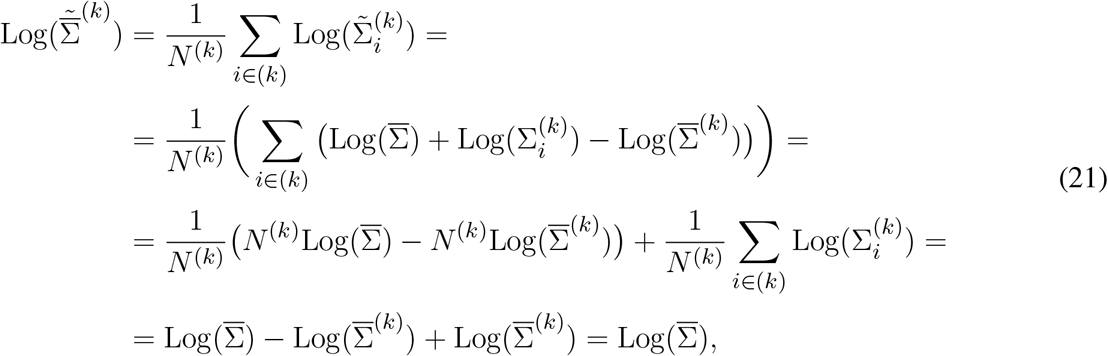

and therefore site means become the global mean 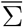. In the case where the term 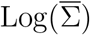 is removed from the transformation rule (14), one gets 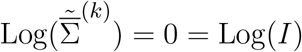, and site means become the identity matrix *I*.

## Notes

https://figshare.com/articles/preprint/Riemannian_geometry_of_functional_connectivity_matrices_for_attention-deficit_hyperactivity_disorder_data_harmonization_ADHD200-derived_functional_connectivity_matrices_for_subjects_in_the_study_results_and_code_/16437534

### Competing Interest Statement

The authors have declared no competing interest.

### Summary of Updates

Version accepted for publication in Frontiers in Neuroinformatics

